# Trustworthy super-resolution reconstruction across spatial omics modalities

**DOI:** 10.64898/2026.09.25.754471

**Authors:** Menghan Li, Xutao Wang, Su Xu, Yuxin Yin, Dong Chen, Xingche Guo, Dongyuan Song

## Abstract

Sequencing-based spatial omics platforms provide scalable and unbiased profiling of transcriptomic, epigenomic, and isoform-resolved signals, but their spatial resolution remains limited because each capture unit aggregates molecular information from multiple cells. Existing computational enhancement methods reconstruct high-resolution spatial gene expression from spot-level data by integrating histology, but most are designed primarily for spatial transcriptomics, rely predominantly on histological features, and lack appropriate validation strategies, leading to overfitting, spurious spatial patterns, and limited reliability. Here we present spEnhance, a generalizable and trustworthy computational framework for super-resolution enhancement of spot-level spatial omics data. spEnhance integrates histological features with complementary molecular information, including single-cell RNA-seq references and gene co-expression structure, to reconstruct high-resolution spatial molecular profiles. To enable reliable model selection from a single tissue section, spEnhance introduces a count-splitting strategy that generates statistically independent training and validation sets from spot-level measurements. spEnhance further quantifies prediction reliability through predictive residuals, providing an interpretable proxy for spatially resolved uncertainty. Comprehensive benchmarking across multiple tissues, platforms, and molecular modalities demonstrates that spEnhance achieves state-of-the-art accuracy, recovers fine-grained tissue structures, mitigates overfitting, and provides calibrated reliability estimates. Beyond spatial transcriptomics, spEnhance extends to isoform-level, epigenomics, proteomic and metabolomic spatial omics. Collectively, spEnhance establishes a modality-general and trustworthy framework for enhancing spatial omics data, enabling more accurate and reliable investigation of spatially resolved molecular regulation.

## 1 Introduction

Understanding how molecular programs are spatially organized within tissues is fundamental to deciphering tissue architecture, defining cellular states, and elucidating their dynamic changes across physiological and pathological processes [1, 2]. Cells exist within spatially heterogeneous environments, where molecular activities vary continuously across space and give rise to complex tissue organization. Accurately resolving such spatial variation at high resolution is therefore essential for linking molecular patterns to tissue architecture and cellular states.

Spatial omics technologies rapidly expanded to profile multiple molecular layers, including transcriptomics, epigenomics, proteomics, and metabolomics. Representative technologies include spatial transcriptomics platforms such as ST [3], Visium [4], Xenium [5], Stereo-seq [6], Slide-seq [7, 8], seqFISH [9], STARmap [10], and MERFISH [11]; spatial epigenomic approaches such as spatial ATAC-seq [12], spatial ATAC-RNA-seq [13], spatial CUT&Tag [14], and SPACE-seq [15]; as well as spatial proteomic and metabolomic methods including DBiT-seq [16], spatial-CITE-seq [17], CODEX [18], and spatial metabolomics technologies [19]. Despite this diversity, most spatial omics platforms can be broadly categorized into imaging-based and sequencing-based approaches. Imaging-based methods achieve subcellular spatial resolution through highly multiplexed fluorescence imaging, but are typically constrained by limited molecular coverage and throughput. In contrast, sequencing-based spatial omics technologies provide unbiased, genome-wide measurements through spatial barcoding strategies, and have become the predominant approach due to their scalability and versatility. Importantly, by directly capturing transcript sequences, these approaches also enable characterization of transcript isoform diversity and post-transcriptional regulation, including alternative splicing (AS) and alternative polyadenylation (APA), particularly when combined with long-read sequencing or event-specific inference methods [20, 21]. However, the effective molecular resolution of sequencing-based approaches remains constrained by an inherent trade-off between spatial granularity and molecular sensitivity. Although recent technologies have achieved single-cell or subcellular-scale spatial profiling, reducing capture size often leads to decreased molecular capture efficiency and increased sparsity, requiring spatial aggregation to obtain sufficient signal. In addition, technical factors such as transcript diffusion, degradation, and capture variability can further compromise spatial fidelity. Consequently, fine-scale molecular heterogeneity and tissue architecture remain challenging to resolve from direct measurements alone.

To address the limited resolution of sequencing-based spatial omics, a variety of computational methods have been developed to reconstruct high-resolution spatial molecular profiles from spot-level measurements by leveraging histological images. Representative approaches, including TESLA [22], iStar [23], scstGCN [24], soscope [25], FineST [26], and iSCALE [27], primarily focus on enhancing spatial transcriptomics data through integration of histology-derived features. While these methods have demonstrated improved spatial resolution and biological interpretability, they share several critical limitations. First, most existing approaches are specifically designed for transcriptomic measurements and are not readily generalizable to other spatial omics modalities, such as spatial epigenomics, proteomics, or isoform-level analyses including AS and APA. Second, most methods lack rigorous validation strategies. Large-scale, independent spatial transcriptomic datasets remain scarce because spatial profiling experiments are costly and relatively low-throughput. Consequently, many approaches are developed and evaluated using data from the same tissue section or from closely related sections, increasing the risk of overfitting and potentially overstating their generalizability. Third, many existing approaches primarily rely on histological features without integrating complementary molecular information, such as single-cell transcriptomic references or biologically informed gene expression programs. This limitation prevents models from leveraging intrinsic molecular relationships, including coordinated gene co-expression patterns, which are essential for accurately capturing biological variability across tissues. Finally, most existing methods do not provide quantitative measures of prediction uncertainty, making it difficult to assess the confidence and robustness of reconstructed spatial signals.

Here, we introduce spEnhance, a generalizable and trustworthy computational framework for super-resolution enhancement of spatial omics data. spEnhance integrates histological features with complementary molecular information, including single-cell reference data and gene co-expression structure, to reconstruct high-resolution spatial molecular profiles. To mitigate overfitting and enable reliable model selection when only a single measured section is available, we leverage count-splitting [28], which partitions the observed count matrix into statistically independent components, allowing one partition to be used for model training and another for validation. This design enables model selection without evaluating candidate models on the same counts used for fitting, thereby reducing information leakage and improving the robustness of model training. Building on this validation framework, spEnhance further leverages held-out count-split measurements to calibrate prediction uncertainty and quantify the reliability of enhanced spatial expression estimates.

Beyond spatial transcriptomics, spEnhance is designed as a modality-general framework that can be readily extended to diverse spatial omics data types, including spatial AS, APA, epigenomics, proteomics, and metabolomics. By operating directly on spatially resolved molecular measurements rather than transcriptomics-specific assumptions, spEnhance enables unified enhancement across molecular layers and facilitates multi-dimensional characterization of spatial gene regulation. Through comprehensive benchmarking across diverse datasets and platforms, we demonstrate that spEnhance achieves improved accuracy, recovers fine-grained spatial structures, and provides calibrated and interpretable predictions. Collectively, these features establish spEnhance as a generalizable and trustworthy framework for enhancing spatial omics data and enabling more reliable investigation of spatially resolved molecular regulation.

## 2 Results

### 2.1 Overview of spEnhance

spEnhance is a trustworthy and modality-general computational framework for super-resolution enhancement of spatial omics data. spEnhance transforms low-resolution spatial molecular measurements into high-resolution spatial profiles by jointly leveraging matched histology, spatial neighborhood structure, single-cell reference information and intrinsic molecular co-variation patterns (Fig. 1a). Unlike approaches that are tailored to a single molecular modality or rely primarily on histological similarity, spEnhance treats spatial omics measurements as a general spatially indexed molecular matrix, making it applicable not only to spatial transcriptomics but also to other spatial omics modalities, including spatial proteomics, spatial chromatin accessibility profiling, spatial metabolomics, spatial chromatin accessibility, and spatial isoform measurements.

**Fig. 1:**
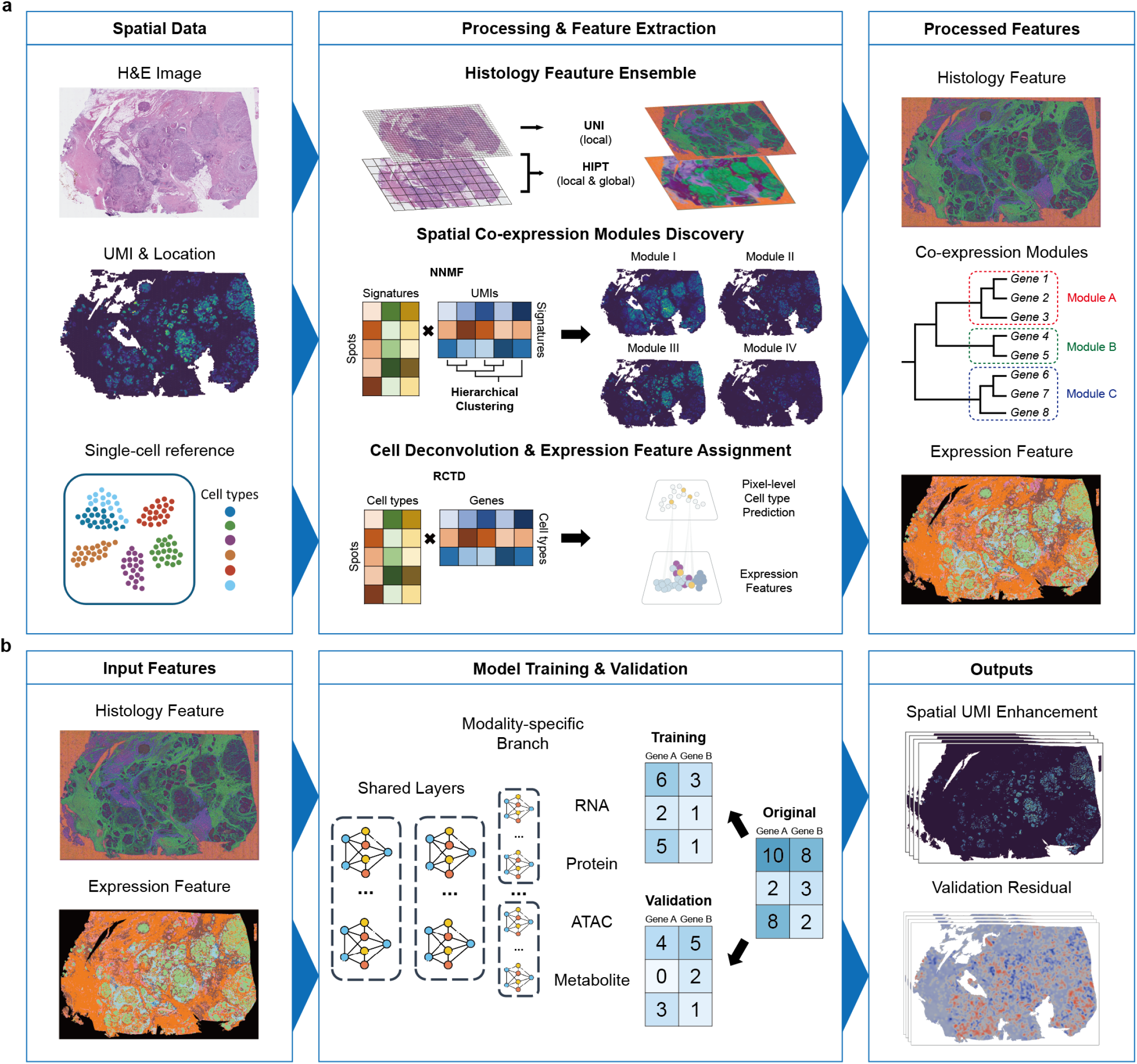
Overview of spEnhance workflow. **a**, spEnhance takes three types of inputs: the H&E image of the tissue section, its spot-level gene expression matrix, and an optional single-cell reference dataset (left). Histology features are extracted using two transformer-based models, and cell-type proportions are inferred via spot deconvolution to generate expression-derived features (middle). Spatial co-expression modules are identified and treated as separate prediction tasks, and a count split strategy produces independent training and validation data. **b**, Leveraging these histology and expression embeddings, spEnhance employs a multi-task learning framework to reconstruct high-resolution spatial gene expression profiles (right). The framework also outputs predictive residuals, which provide a quantitative measure of prediction reliability

spEnhance consists of three core components: a morphology encoder, a reference-and co-expression-guided representation module and a graph-based super-resolution predictor. First, the morphology encoder extracts high-resolution image features from the matched histological image and assigns them to dense spatial locations, thereby capturing local tissue architecture that is not directly resolved by the original spot-level measurements. Second, spEnhance integrates single-cell reference information and gene co-expression structure to learn biologically constrained latent representations. This step provides molecular context beyond local histological appearance and helps the model distinguish spatial domains that may appear morphologically similar but differ in cellular composition or molecular state. Third, the graph-based super-resolution predictor propagates information across neighboring spatial units while preserving local heterogeneity, generating dense molecular maps at pixel-level or cell-level resolution.

A central feature of spEnhance is its emphasis on trustworthy prediction. Because ground-truth high-resolution measurements are unavailable for low-resolution spatial omics datasets, model evaluation and early stopping are challenging in the single-slice enhancement setting. To address this, spEnhance incorporates an internal count-splitting validation strategy that separates the observed molecular counts into training and validation components, enabling prediction performance to be monitored without requiring external high-resolution ground truth. In addition, spEnhance estimates predictive residuals to quantify location- and feature-specific reliability, allowing users to distinguish robustly recovered spatial patterns from uncertain predictions. These design choices make spEnhance not only a super-resolution model, but also a reliability-aware framework for spatial omics enhancement.

The output of spEnhance is a collection of high-resolution spatial molecular maps that can be directly used for downstream biological analysis, including visualization of spatially variable features, identification of fine-scale tissue domains, spatial clustering, cell-type or state mapping and cross-modality comparison. By combining histological information, single-cell reference knowledge, molecular co-variation and graph-based spatial modeling within a unified validation-aware framework, spEnhance enables robust reconstruction of fine-grained spatial molecular organization across diverse tissues, platforms and omics modalities.

### 2.2 spEnhance accurately reconstructs high-resolution spatial expression across diverse benchmark datasets

We systematically benchmarked spEnhance on semi-synthetic spatial transcriptomic datasets derived from human and mouse tissues, spanning multiple organs, disease contexts and measurement platforms. To establish quantitative ground truth, we used Xenium and Xenium Prime datasets, which provide transcript measurements at subcellular resolution. We aggregated the original high-resolution transcript maps according to the spot size, spacing and hexagonal geometry of the Visium platform to generate pseudo-Visium inputs. Models were trained or applied using these pseudo-spot measurements, and their predictions were evaluated against the original high-resolution data.

Across all evaluated datasets, spEnhance consistently outperformed three representative super-resolution baselines, scstGCN, iStar and TESLA, in gene-wise structural similarity index measure (SSIM) and root mean square error (RMSE) (Fig. 2a). The improvement was observed across tissue types and platforms, indicating that spEnhance robustly recovers spatial molecular structure. For example, in the human ovarian carcinoma dataset, spEnhance achieved a median SSIM of 0.9133 and a median RMSE of 0.050, substantially exceeding the performance of competing methods.

**Fig. 2:**
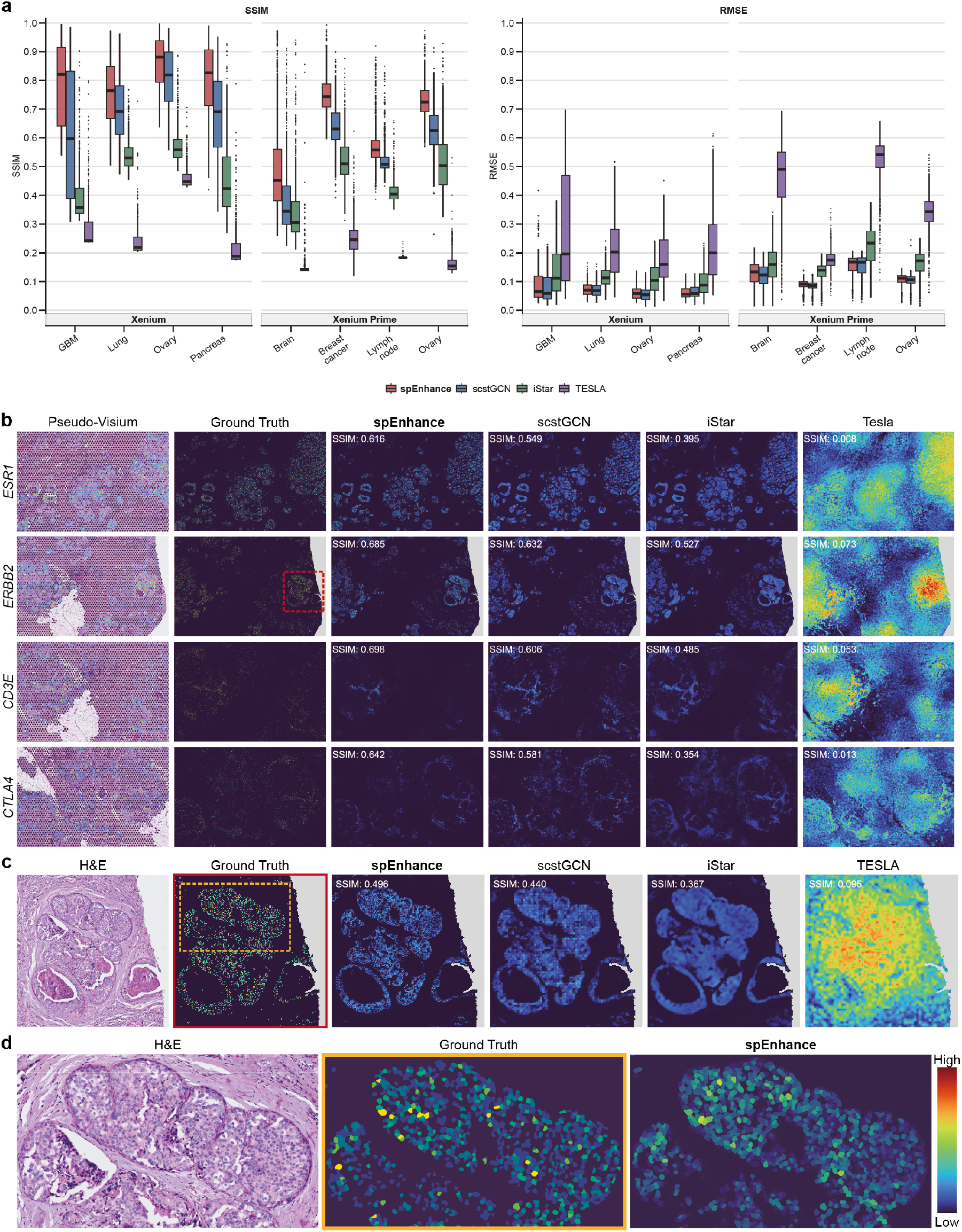
spEnhance recovers high-resolution spatial expression. **a**, Gene-wise comparison in terms of Structural Similarity Index Measure (SSIM, left) and Root Mean Square Error (RMSE, right) across all datasets against scstGCN, iStar, and TESLA. **b**, Visualization of ground truth gene expression measured by Xenium Prime and predicted super-resolution gene expression by spEnhance, scstGCN, iStar, and TESLA. Each column represents a gene. The first two rows correspond to pseudo-Visium data and ground truth revealed by Xenium Prime, respectively. Subsequent rows represent prediction results by each model. **c**, Zoomed-in view of the subregion outlined by the red box in **b**. Histological features, Xenium Prime groundtruth expression, and reconstructed spatial gene expression are shown for the same region. **d**, Visualization of single-cell-level gene expression of a sub-region indicated by the yellow box in **c**. Histology image, Xenium Prime ground truth and predicted gene expression by spEnhance were presented. Pixel-level reconstructed gene expression was aggregated to overlapped cell masks. Nuclei were segmented using Cellpose-SAM.

To examine how these quantitative improvements translated into reconstructed spatial patterns, we examined representative genes in a human breast cancer Xenium Prime dataset. Four genes with distinct biological and spatial patterns were selected: *ERBB2* and *ESR1*, which mark major tumor-associated programmes, and *CD3E* and *CTLA4*, which capture immune-enriched regions. Compared with the pseudo-Visium input and baseline predictions, spEnhance more faithfully recovered the spatial distribution and expression scale of the high-resolution ground truth (Fig. 2b). In contrast, competing methods frequently produced over-smoothed, spatially distorted or scale-shifted reconstructions.

To assess fine-scale spatial fidelity, we further zoomed into an *ERBB2* -enriched subregion. spEnhance preserved sharp tissue boundaries, local gradients and focal expression hotspots that were attenuated or blurred by other approaches (Fig. 2c).

These results show that spEnhance can recover local molecular structures that are obscured by spot-level averaging but are important for interpreting tumor architecture.

Because spEnhance generates dense pixel-level predictions, we further asked whether these reconstructions could support cell-level analysis. We performed nuclei segmentation with Cellpose-SAM, expanded nuclei to approximate cell masks and assigned pixel-level expression to individual cells [29]. The resulting cell-level *ERBB2* expression map closely matched the Xenium Prime ground truth in both spatial localization and cell-to-cell variability (Fig. 2d). Together, these benchmarks demonstrate that spEnhance provides accurate and biologically meaningful super-resolution reconstructions from low-resolution spatial transcriptomic inputs.

### 2.3 spEnhance enables data-driven model selection and reliable uncertainty quantification

A central challenge in training spatial enhancement models is the lack of independent validation data. Unlike conventional machine-learning settings with many exchangeable samples, spatial omics studies often contain only a few tissue sections. Moreover, substantial biological and technical heterogeneity across sections can make an across-section training–validation split infeasible or unrepresentative. Consequently, existing methods commonly use fixed training schedules rather than data-driven stopping criteria. For example, the released implementations of iStar and scstGCN use fixed schedules of 400 and 500 epochs, respectively. Such arbitrary choices increase the risk of overfitting and make prediction reliability difficult to assess. To construct validation data within a single tissue section, a simple approach is to randomly partition spots into training and validation sets, referred as Random Split. However, neighboring spots are spatially correlated, and withholding spots reduces spatial coverage while changing the prediction task from reconstruction at observed locations to interpolation at unobserved locations. We therefore developed spEnhance-CS by adapting count splitting to spatial count data. Count splitting partitions the observed counts at each spot into training and validation components while retaining all spot locations in both datasets. Under the assumed count model, the resulting training and validation counts are independent conditional on the latent expected expression at each spot (Methods).

Figure 3 presents representative results from the Breast Cancer Prime dataset. We first evaluated count splitting using semi-synthetic data with available ground truth. For spEnhance-CS, the validation loss initially decreased, reached a clear minimum at epoch 115, and then plateaued or slightly increased, whereas the training loss continued to decrease (Fig. 3a). By comparison, the random-split validation curve reached a later and shallower minimum at approximately epoch 160 and showed a larger apparent separation from the training loss. Although the absolute losses are not directly comparable because the two strategies define different validation targets, count splitting provided a clearer signal for data-driven early stopping. Additional comparisons of the performance of spEnhance-CS and Random Split across the remaining datasets are provided in Supplementary Figure S1.

**Fig. 3:**
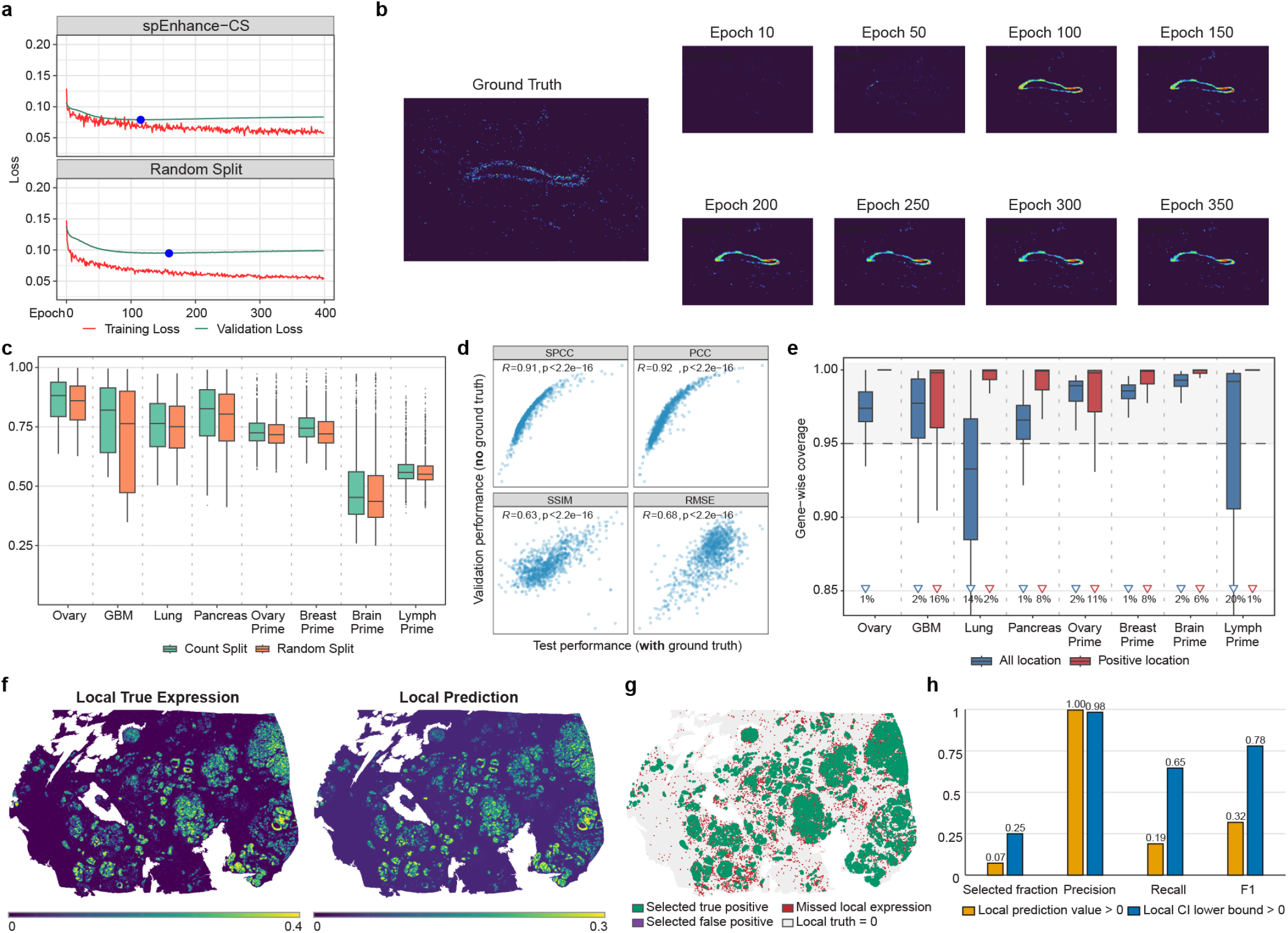
**a**, Training and validation loss curves for spEnhance–Count Split (spEnhance-CS) and a Random Split. The blue star marks the epoch with the minimum validation loss. **b**, Ground-truth gene expression (left) and predicted super-resolution gene expression across training epochs (right). The Structural Similarity Index Measure (SSIM) quantifies the agreement between predictions and ground truth. **c**, Gene-wise SSIM comparing models selected using spEnhance-CS and Random Split across eight benchmark datasets. Boxplots summarize SSIM values across genes within each dataset. **d**, Gene-wise comparison of validation performance (four metrics) based on spEnhance-CS (no ground truth) versus test performance (with ground truth). The Pearson correlation *R* and corresponding *p*-values summarize the concordance between validation-based and true performance across genes.**e**, Gene-wise empirical coverage of nominal 95% local confidence intervals across eight benchmark datasets. Boxplots summarize coverage across unfiltered genes for all tissue locations (blue) and locations with positive local ground-truth expression (red). Downward triangles mark coverage below 0.85; percentages indicate the proportion of genes below this threshold. **f**, Spatial maps of true and predicted local expression for *ESR1* in the Breast Prime dataset. **g**, Classification of tissue locations obtained by thresholding the lower confidence-interval bound at zero. **h**, Comparison of the rounded point-prediction rule and the lower-confidence-bound rule for detecting positive local *ESR1* expression.

The spatial expression pattern of *IL6* further supported the epoch selected by count splitting (Fig. 3b). At epochs 10 and 50, the predicted patterns were incomplete and showed relatively low agreement with the ground truth, with SSIM values of 0.64 and 0.63, respectively. Prediction accuracy improved substantially near the selected epoch, reaching an SSIM of 0.75 at epoch 100. It then remained largely stable, with SSIM values of 0.73–0.74 from epochs 150 to 350, despite the continued decrease in training loss. Across all genes and eight benchmark datasets, models selected using count splitting achieved higher median gene-wise SSIM than those selected using random splitting (Fig. 3c).

Beyond model selection, count-split validation performance provided gene-specific estimates of prediction quality. Genewise validation and ground-truth test performance were strongly correlated for SPCC (*R* = 0.91), PCC (*R* = 0.92), SSIM (*R* = 0.63), and RMSE (*R* = 0.68); all correlations had *p <* 2.2 *×* 10^*−*16^ (Fig. 3d). Thus, validation metrics computed without super-resolution ground truth can reliably distinguish genes with higher and lower reconstruction accuracy.

We next used count-split validation residuals to quantify local prediction uncertainty. Because the validation residuals are observed at the spot level, they do not directly support confidence intervals for individual pixels. We therefore defined the inferential target as a local-neighborhood average: for each target pixel, the local prediction was calculated by averaging the enhanced values within a spot-footprint window centered on that pixel. We then constructed a 95% confidence interval (CI) for this local mean (Methods).

Across unfiltered genes, the local CIs showed broadly reliable coverage in all eight benchmark datasets (Fig. 3e). For evaluation across all tissue locations, the median coverage was at or above the nominal 0.95 level in seven of the eight datasets, with Lung showing lower median coverage. At positive locations, defined as locations with positive local ground-truth expression, median coverage approached 1.0 in every dataset. Some datasets showed broader lower tails in their all-location coverage distributions, particularly Lung and Lymph Prime, indicating gene-and dataset-specific undercoverage. Overall, these results demonstrate that the local CIs are well calibrated for most genes, especially within biologically relevant positive-expression regions (Fig. 3e).

Finally, we used *ESR1* in the Breast Prime dataset to illustrate the practical utility of the local CIs. The local prediction preserved the principal spatial organization of *ESR1* expression observed in the ground truth (Fig. 3f). Thresholding the lower CI bound at zero produced a binary map of local neighborhoods with statistically supported positive expression (Fig. 3g).

Positive local expression was present in 37.8% of the tissue locations. Compared with the rounded point-prediction rule, which selected 7.1% of locations, the lower-bound rule selected 24.8% while maintaining high precision. Specifically, precision decreased only slightly from 0.996 to 0.982, whereas recall increased from 0.188 to 0.645, improving the F1 score from 0.317 to 0.779 (Fig. 3h). The all-location coverage was 0.950, and the median CI width was 0.105. These results show that the local CI substantially improves the detection of truly *ESR1* -positive neighborhoods while retaining high precision.

### 2.4 spEnhance improves tissue segmentation and spatial domain annotation

Having established reconstruction accuracy, we next asked whether the enhanced expression profiles could improve down-stream characterization of tissue architecture and support the downstream domain annotation task. We first analyzed a breast cancer sample for which Visium data from an adjacent section were registered to the H&E image of a Xenium-profiled section. The registered Visium data were enhanced using spEnhance, scstGCN or iStar, followed by spatial clustering with BANKSY [30] (Supplementary Fig. S2). Relative to the pathologist-defined annotations, clustering of the spEnhance reconstruction resolved DCIS-1, DCIS-2 and invasive carcinoma as distinct spatial domains. By contrast, scstGCN failed to recover part of the DCIS-2 region, indicating that incomplete reconstruction of local expression heterogeneity can obscure histologically related but molecularly distinct tumor compartments.

In addition to enhancing individual capture areas, spEnhance can be incorporated into a whole-section reconstruction workflow in which multiple capture areas from the same tissue section are jointly used to predict expression across both profiled and unprofiled regions. We evaluated this setting using a gastric cancer dataset previously analyzed with iSCALE [27]. Across the evaluated genes and spatial regions, spEnhance produced higher structural similarity indices and lower root mean squared errors than iSCALE when compared with Xenium-derived reference measurements (Supplementary Fig. S3). Representative expression maps further illustrated these differences (Supplementary Fig. S3). For *MYH11, TFF2, MKI67* and *CAPN8*, the spatial patterns reconstructed by spEnhance more closely recapitulated the reference distributions. Notably, spEnhance recovered the spatial distribution of *TFF2* in both in-sample regions overlapping the Visium capture areas and out-of-sample regions lacking direct Visium measurements. By comparison, the iSCALE prediction deteriorated substantially outside the sampled regions. Both methods predicted *CAPN8* expression in an out-of-sample region; however, the spEnhance reconstruction exhibited a more fine-grained spatial pattern that was more consistent with both the Xenium reference and the local histological morphology.

The improved expression reconstruction also translated into more accurate tissue segmentation. BANKSY clustering of the spEnhance-enhanced gastric cancer section delineated spatial domains that closely corresponded to pathologist-defined tissue compartments, including tertiary lymphoid structures (TLSs), mucosa and other histologically heterogeneous regions (Fig. 4a). To identify TLSs systematically, we first calculated a spatial TLS marker score using genes associated with lymphoid organization and immune activation[23]. We then incorporated the predictive uncertainty estimated by spEnhance into the composite marker score, allowing regions with consistently high TLS-associated expression to be distinguished from regions in which an apparently high score was weakly supported by the reconstructed data. Spatially contiguous candidate regions were subsequently retained using uncertainty-aware filtering and region-level multiple-testing correction. The resulting TLS calls showed closer agreement with the pathologist annotations than those derived from iSCALE and reduced the false-positive proportion from 35.9% to 17.9% (Fig. 4b,c). Thus, the uncertainty estimates generated by spEnhance can be used not only to evaluate individual gene predictions, but also to improve the reliability of downstream tissue-state annotations derived from combinations of multiple predicted genes.

**Fig. 4:**
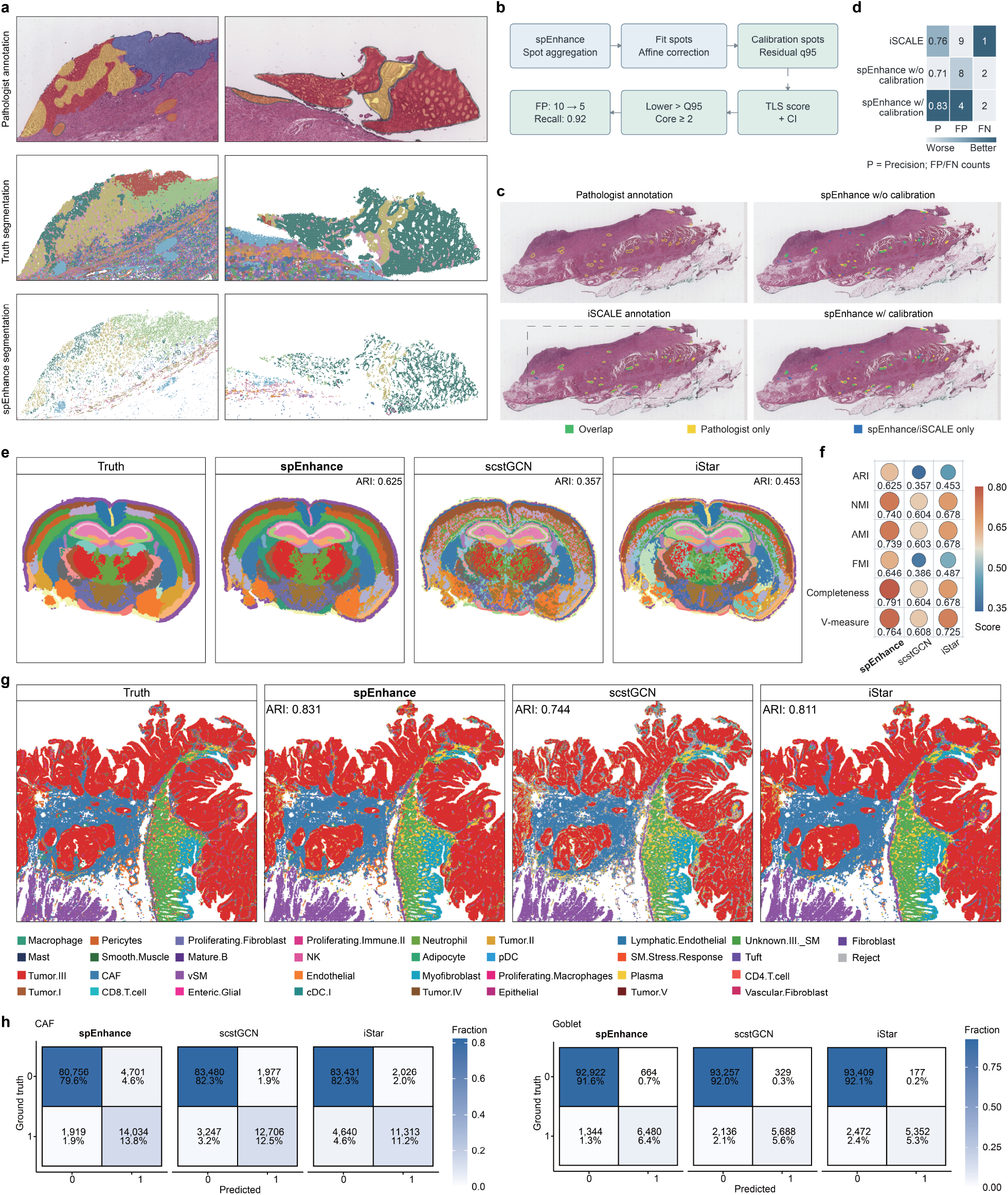
Evaluation of spEnhance tissue segmentation and cell type annotation performance. **a**, Spatial domain segmentation of the human gastric cancer data. Two representative subregions are shown. Pathologist annotation, BANKSY clustering using Xenium ground truth and BANKSY clustering using spEnhance reconstructed spatial expression are presented. Pathologist annotation was adapted from [27]. **b**, Uncertainty-aware workflow for identifying tertiary lymphoid structures (TLSs) from spEnhance-reconstructed expression. **c**, Comparison of pathologist-annotated TLSs with TLSs identified from spEnhance and iSCALE reconstructions. Colors indicate overlapping regions, pathologist-only regions, and methodonly regions. **d**, Precision, false positive, and false negative for iSCALE TLS annotation, spEnhance TLS annotation without calibration and spEnhance TLS annotation with calibration. **e**, BANKSY segmentation of the mouse brain benchmark using ground-truth expression and reconstructions from spEnhance, scstGCN, and iStar. ARI values are shown. **f**, Agreement between reconstructed and ground-truth segmentations across six clustering metrics. g, RCTD cell-type annotations based on ground-truth expression and reconstructions from spEnhance, scstGCN, and iStar. ARI values are shown. **h**, Confusion matrices for representative CAF and goblet-cell annotations.

We then examined whether these improvements generalized across species and spatial transcriptomic platforms. Stereo-seq v2 measurements from a mouse brain section were aggregated to generate pseudo-Visium observations, which were subsequently enhanced using spEnhance, scstGCN or iStar (Supplementary Fig. S4) and segmented with BANKSY. Among the evaluated methods, spEnhance achieved the highest adjusted Rand index relative to the reference segmentation obtained from the original Stereo-seq v2 data (ARI = 0.625; Fig. 4e,f). The spatial domains inferred from the spEnhance reconstruction also exhibited greater anatomical continuity and more closely followed the reference tissue boundaries, whereas the segmentations derived from scstGCN and iStar were comparatively fragmented. Together, these results indicate that spEnhance preserves spatially coherent molecular variation required for tissue segmentation across single-capture-area and whole-section analyses, as well as across imaging-based and sequencing-based spatial transcriptomic platforms.

Finally, we evaluated whether the enhanced expression profiles could improve cell-type annotation at near-cellular resolution using the Human CRC Visium HD dataset. The original Visium HD measurements were first aggregated to generate pseudo-Visium observations, which were then enhanced using spEnhance, scstGCN or iStar. To ensure a consistent resolution for downstream comparison, the expression matrices reconstructed by the three methods were subsequently aggregated into 16 *µ*m bins. Cell-type annotation was performed on each reconstructed dataset using RCTD with the same reference and parameter settings. The resulting annotation maps are shown in Fig. 4g. Relative to the competing methods, annotations derived from the spEnhance reconstruction showed greater spatial continuity and closer agreement with the cell-type distribution inferred from the original Visium HD data, particularly in regions containing spatially localized or less abundant cell populations.

We further quantified annotation performance for cancer-associated fibroblasts (CAFs) and goblet cells, two cell types with distinct spatial distributions in the colorectal tumor microenvironment. For each cell type, the RCTD annotation at each 16 *µ*m bin was converted into a binary presence-absence label and compared with the corresponding ground-truth classification derived from the original Visium HD data. The resulting confusion matrices are presented in Fig. 4h. spEnhance recovered both CAF-and goblet-cell-containing bins with fewer false-positive and false-negative assignments than scstGCN and iStar, indicating improved discrimination between cell-type-positive and cell-type-negative regions. These results demonstrate that the fine-scale molecular variation preserved by spEnhance can improve reference-based cell-type annotation after super-resolution reconstruction, extending its utility from tissue-domain identification to the spatial localization of specific stromal and epithelial cell populations.

### 2.5 spEnhance enables high-resolution reconstruction of spatial proteomic and metabolomic data

We next examined whether spEnhance could be extended beyond spatial transcriptomics to reconstruct spatial protein and metabolite measurements. We first analysed a human tonsil dataset profiled using Visium gene expression and antibody-derived tag (ADT) assays. Representative spatial distributions of *MS4A1* and *VIM* RNA, together with their corresponding protein measurements, showed that spEnhance recovered finer-grained spatial patterns than the spot-level observations and the alternative enhancement methods (Fig. 5a). For RNA reconstruction, we compared spEnhance with scstGCN and iStar, whereas protein reconstruction was compared with iStar because scstGCN did not produce sufficiently stable or spatially interpretable protein maps under the evaluated settings. Across both modalities, the spEnhance reconstructions exhibited sharper spatial boundaries and greater intraregional detail while retaining the overall expression patterns observed at the spot level.

**Fig. 5:**
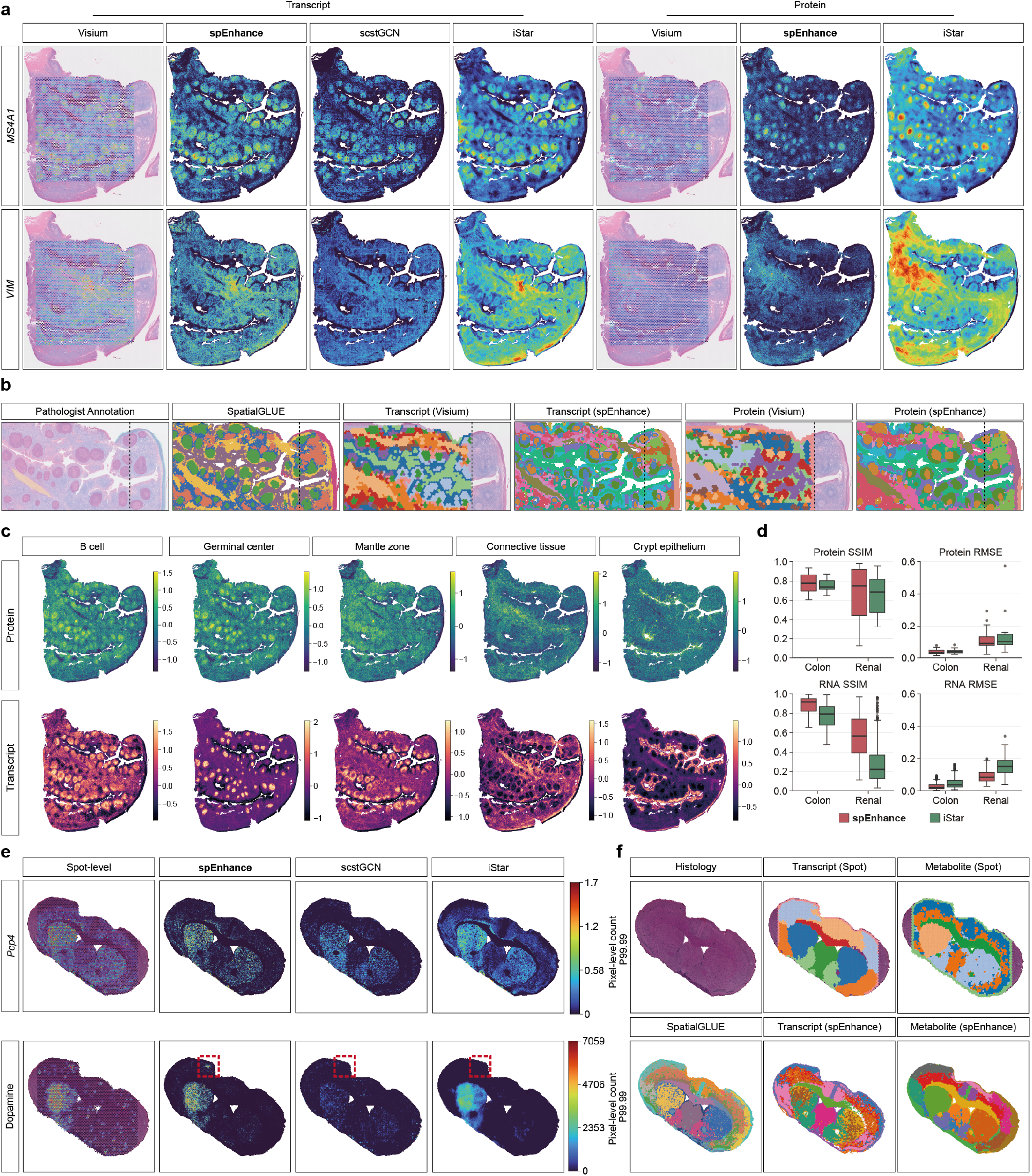
spEnhance extends high-resolution spatial reconstruction to spatial proteomics and metabolomics. **a**, Application of spEnhance to a multimodal Visium human tonsil dataset with paired transcript and protein measurements. Spatial expression patterns of *VIM* and *MS4A1*, together with their corresponding proteins, are shown. **b**, Spatial domain segmentation of the human tonsil dataset. From left to right: pathologist annotation, SpatialGlue clustering based on the enhanced multimodal data, BANKSY clustering using the Visium data, the spEnhance-enhanced transcriptomic data, the Visium proteomic data, and the spEnhance-enhanced proteomic data. Dashed lines indicate the boundary of Visium measurements. **c**, Spatial marker scores for B cell, germinal center, mantle zone, connective tissue, and crypt epithelium compartments, calculated from the spEnhance-enhanced transcriptomic and proteomic profiles, respectively. **d**, Benchmarking of spatial reconstruction accuracy on colon cancer and renal cancer datasets. Reconstruction performance was evaluated using structural similarity index measure (SSIM) and root mean squared error (RMSE), with distributions summarized as boxplots. **e**, Application of spEnhance to spatial metabolomics data. Spatial distributions of *Pcp4* expression and dopamine abundance are shown at the original spot level and after enhancement. Boxes indicate representative subregions highlighted for comparison. **f**, Spatial segmentation of the metabolomics dataset. Histology image, segmentation based on spot-level transcriptomic data, spot-level metabolite data, spEnhance-reconstructed transcriptomic data, spEnhance-reconstructed metabolite data, and SpatialGlue clustering based on the enhanced multimodal profiles were shown.

Having confirmed that spEnhance could reconstruct both RNA and protein signals, we asked whether the enhanced measurements could better delineate functionally distinct tonsillar compartments. Within a representative subregion, we compared the pathologist-defined annotations with modality-specific clustering of the original spot-level measurements, clustering after enhancement with spEnhance, and joint RNA-protein integration using SpatialGlue [31] (Fig. 5b). Clustering of original spot-level transcript or protein measurements did not consistently recover all germinal centers annotated by pathologists and frequently merged germinal center-associated regions with neighboring tissue compartments. The resulting domains were also spatially coarse, reflecting the limited resolution of the original Visium measurements. In contrast, clustering of the spEnhance reconstructions recovered each annotated germinal center in the displayed region and further distinguished the germinal center from its surrounding mantle zone. Joint analysis of the enhanced RNA and protein modalities produced spatial domains that more closely followed the local histological organization and resolved tissue structures at a substantially finer spatial scale.

To characterize these compartments independently of unsupervised clustering, we calculated modality-specific marker scores for major tonsillar cell populations and tissue regions, including B-cell-rich areas, germinal centers, mantle zones, connective tissue and crypt epithelium (Fig. 5c). Marker scores derived from either RNA or protein measurements broadly recapitulated the expected anatomical distributions. The RNA-derived maps generally exhibited sharper boundaries and finer spatial definition, whereas the protein-derived maps showed greater local variability. Although the two modalities were concordant at the level of major tissue compartments, localized differences between RNA-and protein-derived scores were also apparent, suggesting that their spatial distributions were not uniformly coupled across the tissue.

To quantitatively validate protein reconstruction against high-resolution ground truth, we next analyzed a colon cancer dataset containing subcellular-resolution RNA and protein measurements generated with the G4X Spatial Sequencer. We aggregated the original measurements to generate pseudo-Visium observations and used the unaggregated data as the reference for benchmarking. Across both RNA and protein measurements, spEnhance achieved lower root mean squared errors and higher structural similarity indices than iStar (Fig. 5d). These results indicate that the framework can reconstruct spatial protein abundance without requiring modality-specific changes to the overall enhancement strategy.

We next extended this analysis to spatial metabolomics using a paired transcriptomic–metabolomic dataset from a mouse model of Parkinson’s disease, in which the analyzed brain section contained an intact region on one side and a lesioned region on the other. Representative maps of *Pcp4* RNA and dopamine showed that spEnhance more closely preserved the spatial distributions of both transcript and metabolite signals than scstGCN or iStar (Fig. 5e). In particular, a localized dopamine-positive region (red box) that was evident in the original spot-level measurements was recovered by spEnhance but was not reconstructed by either scstGCN or iStar, resulting in an apparent false-negative region in the latter predictions. Thus, spEnhance retained spatially restricted metabolite signals that could be lost during reconstruction by competing approaches. Consistent with these expression-level improvements, spatial clustering based on the enhanced transcriptomic or metabolomic measurements produced domains that more closely corresponded to recognizable neuroanatomical subdivisions than clustering based on the original spot-level measurements (Fig. 5f). Joint integration of the enhanced modalities using SpatialGlue further resolved spatially coherent domains that followed the anatomical organization visible in the H&E image. Together, these analyses demonstrate that spEnhance can be applied across spatial RNA, protein and metabolite measurements and that the resulting high-resolution reconstructions improve both modality-specific characterization and integrative spatial domain analysis.

### 2.6 spEnhance reconstructs spatial isoform and APA patterns at high resolution

We next asked whether spEnhance could extend beyond gene-level expression to resolve transcript-level variation, including alternative polyadenylation (APA) and transcript isoforms, while preserving their quantitative relationships with the corresponding gene-level expression. We therefore added a consistency term to the training objective that constrains the reconstructed expression of individual isoforms or APA sites to remain consistent with the total expression of the corresponding gene.

We first evaluated spEnhance on spatial APA profiles derived from the Visium mouse brain sagittal anterior dataset. Spatial APA expression was inferred from the original Visium data using Infernape [21], and the resulting spot-level APA profiles were subsequently enhanced with spEnhance. Visual comparison of a representative gene, *Dclk1*, showed that spEnhance reconstructed fine-scale expression patterns of its major APA sites while preserving their characteristic spatial distributions observed in the original Visium measurements (Fig. 6a). Compared with reconstructions generated by scstGCN and iStar, the spEnhance results retained spatially coherent APA patterns at substantially increased resolution.

**Fig. 6:**
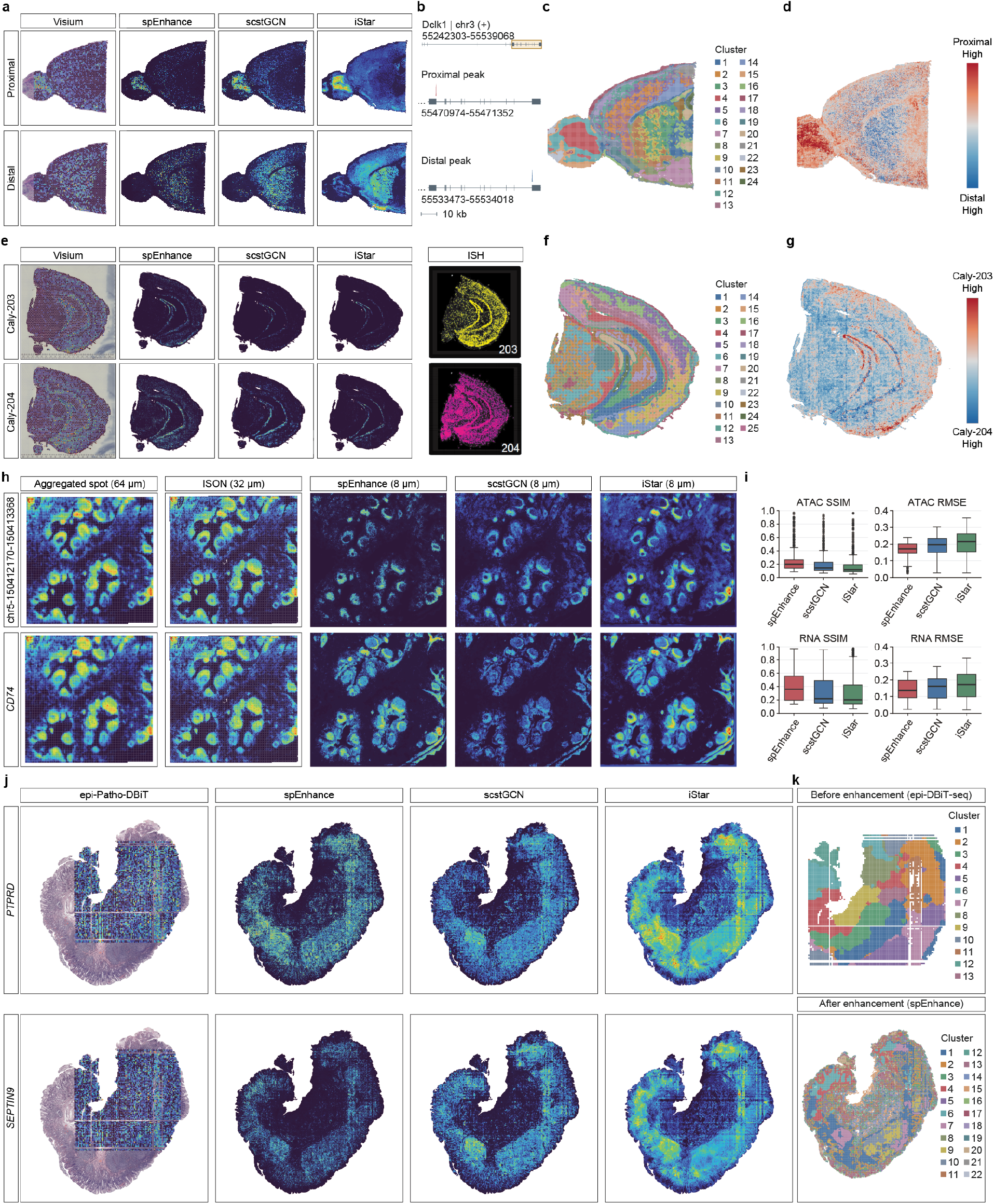
spEnhance resolves high-resolution patterns of alternative polyadenylation, transcript isoform usage, and epigenomic activity. **a–d**, Spatial alternative polyadenylation (APA) analysis of a sagittal mouse brain Visium dataset. APA isoforms were inferred using Infernape and enhanced with spEnhance. **a**, Observed and reconstructed spatial distributions of the proximal and distal *Dclk1* isoforms. **b**, *Dclk1* gene structure showing the two analyzed APA sites. **c**, BANKSY domains inferred from spEnhance-enhanced gene expression. **d**, Pixel-level relative usage of the two *Dclk1* isoforms. **e–g**, Transcript isoform analysis of a coronal mouse brain spatial long-read sequencing dataset. **e**, Observed and reconstructed expression of *Caly-203* and *Caly-204*, with in situ hybridization (ISH) patterns as spatial references. **f**, BANKSY domains inferred from spEnhance-enhanced gene expression. **g**, Pixel-level relative usage of *Caly-203* and *Caly-204*. **h**,**i**, Joint enhancement of spatial transcriptomic and simulated epigenomic profiles from a human tonsil Visium HD dataset. ISON was used to simulate epigenomic measurements at 32 *µ*m resolution, which were aggregated to 64 *µ*m bins for benchmarking. **h**, Representative reconstructions of *CD74* expression and its corresponding ATAC-seq peak. **i**, SSIM and RMSE comparisons across methods. **j**,**k**, Enhancement of FFPE spatial ATAC-seq data from a MALT sample profiled by epi-Patho-DBiT. **j**, Observed and reconstructed activity scores for *SEPTIN9* and *PTPRD*. **k**, BANKSY clustering based on observed epi-Patho-DBiT gene activity scores and those reconstructed by spEnhance.

We next asked whether the enhanced profiles preserved regional variation in relative APA usage. We performed tissue segmentation using the spEnhance-reconstructed gene-level expression profiles. The resulting spatial domains recapitulated major anatomical structures of the mouse brain section (Fig. 6b). We then quantified the relative usage of two dominant *Dclk1* APA sites across the reconstructed section. Distinct regional preferences for the two APA sites remained apparent after enhancement, demonstrating that spEnhance preserved and resolved spatial variation in APA usage at the pixel level (Fig. 6c).

We further evaluated spEnhance using a spatial long-read Visium dataset from a mouse brain coronal section, which provides direct measurements of transcript isoforms. We reconstructed the spatial expression of individual isoforms using spEnhance, scstGCN, and iStar. For the representative gene *Caly*, the reconstructed distributions of isoforms *Caly* -203 and *Caly* -204 showed distinct spatial patterns (Fig. 6d). Notably, the patterns recovered by spEnhance were concordant with the corresponding spatial distributions observed by in situ fluorescence imaging (Fig. 6e), providing orthogonal support for the reconstructed isoform-specific signals. Tissue segmentation based on the reconstructed gene-level expression further recovered spatially organized anatomical domains (Fig. 6f), while analysis of isoform usage revealed region-specific differences between the two *Caly* isoforms at enhanced spatial resolution (Fig. 6g).

Together, these results demonstrate that spEnhance can extend spatial expression reconstruction beyond the gene level to APA sites and transcript isoforms, while preserving both their underlying spatial distributions and their relative usage within the corresponding gene.

### 2.7 spEnhance extends to spatial epigenomic reconstruction

Having demonstrated spEnhance across transcriptomic, proteomic, metabolomic and isoform-level measurements, we finally evaluated its applicability to spatial epigenomic data, for example, spatial ATAC-seq. Because currently available spatial ATAC-seq datasets generally lack paired high-resolution histology images suitable for controlled benchmarking, we first constructed a spatial ATAC-seq benchmark based on the Visium HD human tonsil dataset. We used ISON together with a human tonsil single-cell multiome reference to infer peak-level chromatin accessibility for each spatial bin. To improve the robustness of the inferred accessibility profiles, the original Visium HD measurements were first aggregated to 32 *µ*m bins for ISON inference, and the resulting spatial ATAC profiles were subsequently aggregated to 64 *µ*m resolution as input for spatial enhancement (Fig. 6h, left).

We then applied spEnhance to reconstruct peak-level chromatin accessibility at higher spatial resolution. Representative accessibility maps showed that spEnhance recovered fine-scale spatial patterns from the low-resolution input while preserving the broader spatial organization of individual ATAC peaks (Fig. 6h, right). We compared these reconstructions with those generated by iStar and quantitatively evaluated reconstruction accuracy using structural similarity index measure (SSIM) and root mean squared error (RMSE). Across the benchmark, spEnhance achieved higher SSIM and lower RMSE than iStar, indicating improved recovery of both spatial structure and quantitative accessibility signals (Fig. 6i).

To further assess the applicability of spEnhance to experimentally measured spatial epigenomic data, we applied the method to a human cancer dataset generated using the epi-PathoDBiT-seq platform. spEnhance successfully reconstructed high-resolution spatial chromatin-accessibility patterns from the original measurements, recovering spatially coherent peak-level signals across the tissue section (Fig. 6j). Together, these results demonstrate that spEnhance can be extended from spatial transcriptomic reconstruction to spatial epigenomic data and can recover high-resolution chromatin-accessibility patterns across both computationally generated benchmarks and experimentally measured spatial ATAC-seq datasets.

## 3 Discussion

In this study, we developed spEnhance, a general framework for reconstructing spatial molecular profiles at higher resolution by integrating measured spatial signals with histological features and single-cell references. We systematically evaluated its performance across multiple spatial transcriptomics platforms, tissue types and benchmark settings, and showed that spEnhance can recover fine-scale spatial expression patterns and improve downstream analyses such as tissue segmentation and cell-type annotation. Beyond gene-level transcriptomics, we further extended the framework to other molecular modalities, including chromatin accessibility, proteins, metabolites and transcript isoforms, demonstrating that our strategy can be generalized to spatial data with distinct dimensionality, sparsity and measurement characteristics. Together, these results establish spEnhance as a flexible framework for spatial molecular reconstruction across diverse technologies and data modalities.

A central consideration in spatial enhancement is how to evaluate model performance while an independent high-resolution ground truth is unavailable for the same tissue. In spEnhance, we address this issue by constructing separate training and validation observations through count splitting. Rather than withholding spatial locations or genes, which would alter the spatial or molecular structure of the data, individual molecular counts are stochastically partitioned into conditionally independent subsets. The training split is used for model optimization, whereas the held-out split provides an internal validation signal for model selection and assessment. This strategy therefore enables validation against held-out molecular counts from the same biological specimen, allowing model predictions to be evaluated using measurements that were not used for fitting.

Importantly, spEnhance also provides a multimodal reconstruction framework. Spatial molecular technologies increasingly measure diverse molecular layers, and the characteristics of these data differ substantially in dimensionality, sparsity, dynamic range and measurement noise. We therefore extended the framework to additional modalities by adapting the input representation and learning objective to the statistical properties of each data type while retaining the same general spatial reconstruction strategy. For spatial ATAC-seq, chromatin accessibility can be represented as counts over genomic peaks, resulting in a substantially higher-dimensional and sparser feature space than conventional transcriptomic measurements. Treating individual accessible regions as molecular features enables spEnhance to reconstruct spatial accessibility landscapes at the peak level, from which regulatory activity at genes or genomic regions can subsequently be derived.

Protein and metabolite measurements present a different regime. These assays generally contain substantially fewer molecular features than transcriptome-wide measurements and may exhibit modality-specific abundance distributions and noise characteristics. Rather than assuming that the loss function optimized for RNA counts is universally appropriate, spEnhance allows the reconstruction objective to be adapted to the corresponding modality. This enables the framework to incorporate spatial proteomic or metabolomic measurements while preserving their quantitative structure and leveraging the same histological and spatial information used for transcriptomic reconstruction. Such flexibility may become increasingly important as spatial multi-omic platforms begin to measure multiple molecular layers within the same tissue section.

The framework can also be applied to transcript features below the gene level, including alternative polyadenylation and alternatively spliced isoforms. In contrast to gene-level expression, these measurements are often substantially more sparse because reads assigned to a gene are distributed among multiple transcript or processing states. We therefore treat individual isoforms or alternative processing events as distinct molecular features during reconstruction, allowing their spatial distributions to be recovered without collapsing them into total gene expression. This provides an opportunity to examine spatial variation in transcript usage that may be obscured at the gene level. Importantly, reconstruction of individual isoforms also enables downstream calculation of relative isoform usage at higher spatial resolution, thereby separating changes in transcript processing from changes in overall gene abundance.

Together, these extensions suggest that spatial enhancement can be formulated as a general molecular reconstruction problem rather than a task specific to gene expression. The combination of internal validation through count splitting and modality-aware reconstruction provides a framework for adapting spEnhance to spatial measurements with substantially different statistical properties. As spatial technologies continue to expand from transcriptomics toward chromatin, proteins, metabolites and transcript isoforms, such modality-flexible approaches may facilitate integrated analysis at spatial resolutions beyond those directly provided by the original assays. At the same time, reconstruction cannot substitute for direct measurement, and predictions at enhanced resolution should be interpreted as model-supported estimates rather than independently observed molecular events. Orthogonal experimental validation will therefore remain important, particularly when enhanced maps are used to infer fine-scale regulatory or cellular mechanisms.

## 4 Methods

### 4.1 The algorithm of spEnhance

spEnhance reconstructs high-resolution spatial expression from spot-level spatial transcriptomics or other types of omics techniques by leveraging H&E-stained histology images and single-cell reference information. In the methods section, we use spatial transcriptomics as the primary illustrative example.

### 4.2 Histology feature extraction via dual-model embedding

To enable consistent analysis across whole-slide histological images acquired at varying resolutions, we first rescale each raw slide so that one pixel corresponds to a physical area of 0.5 *×* 0.5 *µ*m^2^. Under this calibration, a 16 *×*16 pixel tile spans an 8*×* 8 *µ*m^2^region (approximately the size of a single cell). Suppose the histology image has height *H* and width *W* (in pixels). We then extract histological features via a dual-model strategy: **(1) HIPT** [32]: A hierarchical vision transformer that captures multi-scale tissue architecture. **(2) UNI** [33]: A universal pathology foundation model trained on over 100 million H&E tiles collected from diverse tissue types and disease contexts. Combining two complementary embeddings allows us to combine global contextual representations (HIPT) with fine-grained local morphology (HIPT & UNI). A unified histology embedding is constructed by applying principal component analysis (PCA) to each model’s features (retaining components that explain at least 99% of the variance) and then concatenating the reduced embeddings. Let 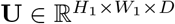denote the histology embedding.

#### Global histological feature extraction using HIPT

In our implementation, we adopted the pretrained model named **HIPT** in Chen et al.[32]. HIPT is a hierarchical vision transformer designed for histopathological image analysis. The preprocessed H&E image was hierarchically divided into large tiles and small tiles, where each small tile is a subregion within a large tile. Large tiles encode global-level features, while small tiles capture local-level features. Let **X** *∈* ℝ^*H×W×*3^ be the RGB-channel H&E image with height *H* and width *W*. Then the downsized image of size *H*_1_ = *H/*16 and *W*_1_ = *W/*16 corresponds to approximately single-cell resolution. We first divide **X** into a (*H/*256) *×* (*W/*256) grid of 256*×*256-pixel tiles: 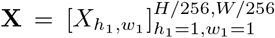, where 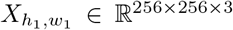. Next, each 256*×*256-pixel tile is partitioned into non-overlapping 16*×*16 subtiles each with 16*×*16-pixel: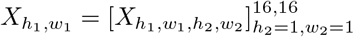, where 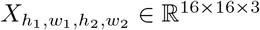.

To capture multi-scale morphological patterns, we adopt a hierarchical vision transformer (HViT) composed of a local vision transformer (ViT) *F*_*L*_ and a global ViT *F*_*G*_. For the local ViT, we partition a 256*×*256 tile indexed by (*h*_1_, *w*_1_) into non-overlapping 16*×*16 subtiles and map each subtile to a *C*_1_-dimensional patch embedding:

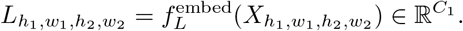

Next, we aggregate the 16×16 patch-level embeddings within that tile and map it into a *C*_2_-dimensional region embedding using a Multi-head Self-Attention (MSA) layer

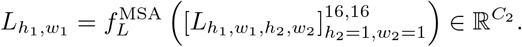

As a result, 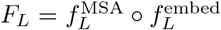. The global ViT maps the grid of slide-level local features for the entire image to a grid of slide-level global features of the same dimension:

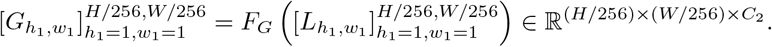

Note that both the local and global ViTs operate on flattened token sequences; the global ViT then reshapes (unflattens) its outputs back to a grid.

Finally, we concatenate the patch-level features 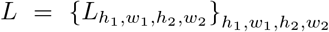 (ordered consistently across all subtiles/tiles), the slide-level features 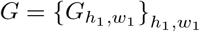, and the RGB channels of the original image *X*. Specifically, we apply bicubic interpolation on *G* to match the spatial resolution of *L* (i.e., *H*_1_ *× W*_1_) and downsample *X* to the same resolution via average pooling. Concatenating along the channel axis yields 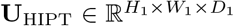 with *D*_1_ = *C*_1_ + *C*_2_ + 3. HIPT adopted a DINO[34] framework to train the model on The Cancer Genome Atlas data. We found the pretrained model was capable of capturing histology features well enough and thus skipped the fine-tuning step. Our implementation was based primarily on the pipeline in [23].

#### Fine-grained histological feature extraction using UNI

To obtain fine-grained histological features, we divide image **X** into a (*H/*224) *×* (*W/*224)-grid of 224*×*224-pixel tiles: 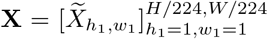 where 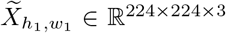. We extract histology feature using another ViT *F*_1_. Each 224*×*224-pixel tile 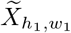 was mapped to a fine-grained representation and then reshaped into a tensor of shape 14 *×* 14 *× C*_3_, that is,

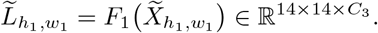

Here we adopted a pretrained model named **UNI** [33], which is a universal pathology foundation model trained on over 100 million H&E tiles collected from diverse tissue types and disease contexts. It adopts a DINOv2 [35] framework to learn generalizable histological representations, enabling robust transfer across tasks without fine-tuning.

To complement the learned features, we augment the UNI features 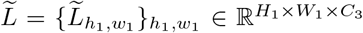 by concatenating,along the channel dimension, (i) the downsampled RGB image (3 channels) and (ii) two positional channels encoding the normalized *x*-and *y*-coordinates (in [0, 1]). These auxiliary inputs provide spatial context and preserve tissue geometry. The resulting tensor is 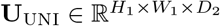 with *D*_2_ = *C*_3_ + 3 + 2.

### 4.3 Single-cell reference guided gene expression embedding

A single-cell reference provides complementary molecular context beyond histology and can improve prediction. Below, we describe how we construct a gene-expression embedding from the single-cell reference.

#### Cell-type deconvolution using RCTD [36]

We built a single-cell reference by performing standard quality control and normalization, clustering cells, and manually curating cluster identities to obtain reliable cell-type labels. Next, we computed the cell type–by–gene reference matrix **R** ℝ^*C×G*^, where each entry *R*_*c,g*_ is the mean expression of gene *g* across all cells of type *c*. Using the Visium count matrix 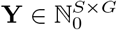 we then applied RCTD to deconvolve each spot, yielding **Π** = [*π*_*s,c*_] ∈ * [0, 1]^*S×C*^, which estimates the fraction of each cell type present in spot *s* = 1, … , *S* (rows summing to one).

#### Pixel-level cell type prediction

To predict pixel-level cell types, we train a graph convolutional network (GCN) [37] that maps the histology embedding **U** to pixel-wise probabilities 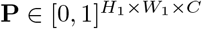, using the spot-level deconvolution **Π** for weak supervision. In training, we first associate pixels with their corresponding spots. For pixel *v* = (*h, w*), let 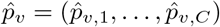 denote the predicted cell-type probabilities. Averaging over pixels assigned to spot *s* yields 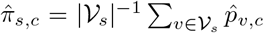, where *V*_*s*_ is the collection of pixels covered by spot *s*. We train the model by minimizing the Kullback–Leibler divergence from the spot-level target *π*_*s*_ = (*π*_*s*,1_, … , *π*_*s,C*_) to the aggregated prediction 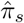. For pixel-level evaluation, we partition the *H*_1_ *× W*_1_ pixel grids into square regions, each of which covers the spatial extent of one spot. The partitioning was performed with a stride parameter *k*, such that adjacent regions overlapped by *k* rows or columns. Within each region, we apply the trained GCN to produce pixel-level predictions. For pixel *v* in overlapping regions, the final predicted probability vector was the average over all regions.

#### Gene expression feature assignment

Given the cell type-by gene reference matrix **R** and the pixel-level cell-type probabilities **P**, we assign a gene-expression value at pixel 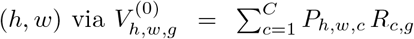. Collecting all genes and pixels forms the tensor 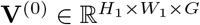. To extract compact representations, we reduce the gene dimension using truncated SVD to obtain the gene feature embedding 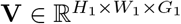.

### 4.4 Trustworthy prediction of super-resolution spatial gene expression

#### The predictive model

To predict pixel-level spatial gene expression 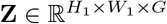, we train a deep neural network that maps the fused histology–gene embeddings 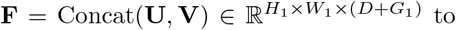 **Z**, using the spot-level count matrix **Y** for weak supervision. Our model is a graph convolutional network (GCN) with two shared graph-convolution layers (512 hidden units each). To reflect the tendency of co-expressed genes to exhibit similar behavior, we adopt a **multi-task learning** design [38] in which genes are grouped into *M* spatial co-expression modules, identified via non-negative matrix factorization (NNMF [39]) on the count matrix **Y** followed by hierarchical clustering. After the shared layers, each module is predicted by its own head (a linear layer mapping 512 to the number of genes in the module). A dropout layer (*p* = 0.5) is applied between the shared representation and each module-specific head to reduce overfitting.

#### Validation set construction

To address the common lack of replicates in spatial transcriptomics, where the entire spot evel dataset is often used for training without a hold-out set, we adopt a **count splitting** approach [40]: Given a spot-level count matrix 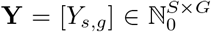, we partition each entry as 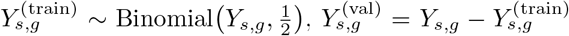. Under a Poisson model assumption, count splitting ensures that 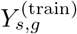 and 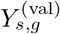 follow the same distribution, and are conditionally independent given their mean value. This construction therefore yields statistically independent training and validation sets suitable for unbiased assessment of predictive performance.

#### Trustworthy model training

For training, pixel-level predictions are aggregated to the spot level via the predefined pixel–spot mapping *V*_*s*_. The network is fit by minimizing the square error loss between aggregated predictions and observed spot counts:

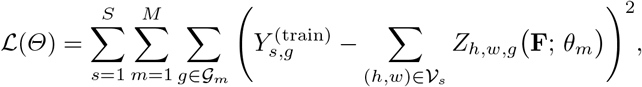

where 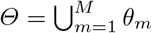 collects the module-specific parameters and *G*_*m*_ denotes the set of genes in module *m*. After each epoch, the validation loss is computed by replacing **Y**^(val)^ in *L* with **Y**^(val)^. The model checkpoint with the lowest validation loss is retained as the final trained model.

#### Model evaluation and predictive performance quantification

For pixel-level prediction, the trained model was applied to the complete (unsplit) embedding set **F**, yielding the final super-resolution estimates of spatial gene expression. To quantify predictive performance, we first aggregate pixel-level predictions to the spot level and then compute spot-level predictive residuals between the observed and predicted spot-level counts. To stabilize the variance for count data, spEnhance uses square root transformation or Pearson residuals. This residual provides a convenient metric for assessing discrepancies between observed and predicted spot-level counts: positive values indicate underestimation, negative values indicate overestimation, and values near zero correspond to well-calibrated predictions.

#### Trustworthy quantification of uncertainty

Uncertainty was quantified post hoc once the predictions at pixel level were obtained. The spot-level predictions for gene *g* at spot *s* were computed as:

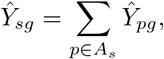

where *Ŷ_pg_* denotes the predicted expression of gene *g* at pixel *p*, and *A*_*s*_ is the set of pixels contributing to spot *s*. The aggregated predictions were then compared with the 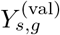. Both observed and predicted spot counts were transformed using the prespecified Anscombe-type variance-stabilizing transformation

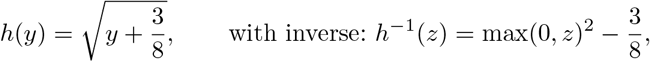

followed by truncation at zero. For each gene *g*, absolute calibration residuals were calculated as

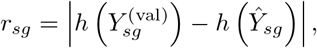

At a nominal error level *α* = 0.05, the gene-specific conformal error bound was defined as

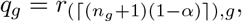

where *r*_(*k*),*g*_ is the *k*-th ordered calibration residual for gene *g* and *n*_*g*_ is the number of usable count-split validation spots for gene *g*.

Because calibration was performed using observed spot-level counts, the primary inferential target was local-average expression around a pixel, with

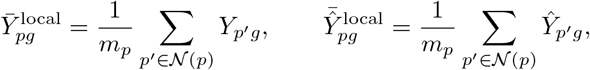

where *N* (*p*) is the local neighborhood centered at pixel *p*, and *m*_*p*_ = | *N* (*p*) |. For benchmark datasets, *N* (*p*) is the same equal-weight square footprint used to aggregate pixels back to spots. If radius is *R* in the raw image coordinate system (*x, y*), and the prediction-grid scale factor is *r*, then

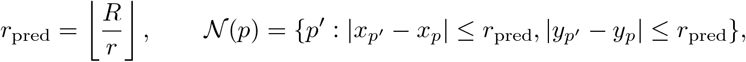

For a neighborhood *N* (*p*) centered at pixel *p*, defined using the same spatial footprint employed for spot aggregation, the 100(1 *− α*)% uncertainty interval for local-average expression was then calculated as

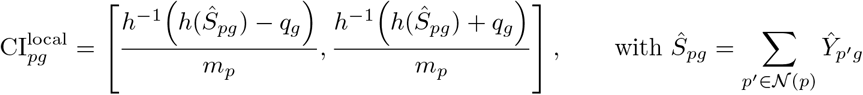

For datasets with pixel truth, all-tissue local-average coverage for gene *g* and across total number of pixels *M* is

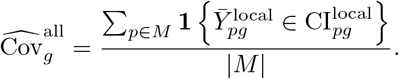

The expressed local target is defined as

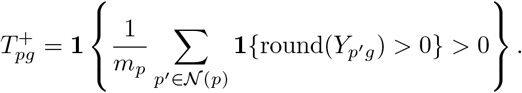

In words, a local neighborhood is truth-positive if at least one rounded-positive true pixel appears within the local neighborhood. The expressed tissue local-average coverage is

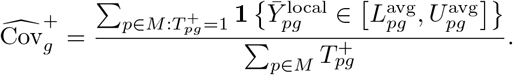

Two binary decision rules were compared. The rounded local prediction rule is

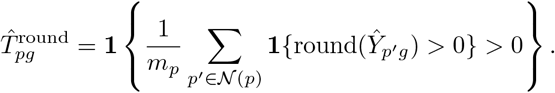

The high-confidence lower-bound rule is

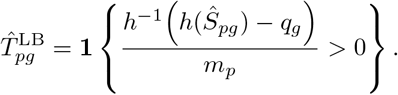

Precision, recall, and F1 are evaluated against 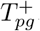. The lower-bound rule is designed to select spatial regions where positive local expression is statistically supported, rather than merely where the point prediction is nonzero.

### 4.5 Joint enhancement of RNA and complementary molecular modalities

#### Multi-modal prediction architecture

To extend the super-resolution framework to spatial ATAC-seq, protein, and metabolite measurements, we adopt a two-branch multi-task architecture. Given the fused spatial embedding 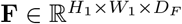, a shared spatial encoder first produces a common pixel-level representation. The representation is subsequently passed to an RNA branch and a modality-specific branch:

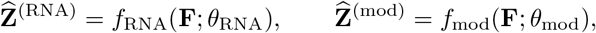

where 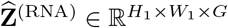 denotes pixel-level gene expression and 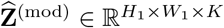 denotes the enhanced complementary modality. Here, the *K* modality-specific features correspond to chromatin-accessibility peaks for ATAC-seq, proteins for spatial proteomics, or metabolites for spatial metabolomics. The two prediction branches have separate output parameters but are trained jointly with the shared spatial representation, allowing information from RNA and the complementary modality to mutually constrain the learned spatial features.

#### Joint weakly supervised training

Let **Y**^(RNA)^ ∈ ℝ^*S×G*^ and **Y**^(mod)^ ℝ^*S×K*^ denote the spot-level RNA and complementary-modality measurements, respectively. Pixel-level predictions are mapped back to the observed resolution using the same pixel–spot mapping *V*_*s*_:

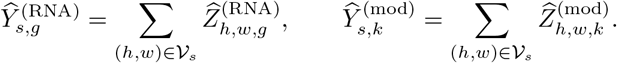

The RNA branch is optimized using the squared-error objective described above:

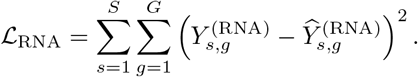

Because ATAC, protein, and metabolite measurements may contain large feature-specific values and technical outliers, we use the Huber loss for the complementary-modality branch:

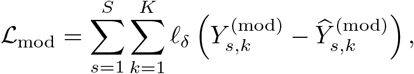

where

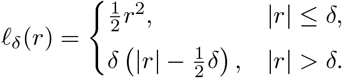

The Huber objective retains quadratic sensitivity to small residuals while reducing the influence of extreme observations. The complete network is trained end-to-end by minimizing

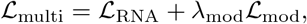

where *λ*_mod_ controls the contribution of the complementary modality and accounts for differences in measurement scale and feature dimensionality. When count splitting is applicable, the split training measurements are used in the joint objective, while the corresponding held-out measurements are used for model selection. After training, the two branches simultaneously produce spatially aligned pixel-level RNA and modality-specific molecular maps.

### 4.5 Consistency-guided enhancement of alternative polyadenylation and alternative splicing

#### Gene- and isoform-level prediction

For alternative polyadenylation (APA) and alternative splicing (AS), the model jointly reconstructs total gene expression and the expression of the corresponding isoforms. For APA, an isoform-level feature represents a transcript or polyadenylation-site-specific product; for AS, it represents a transcript or splice isoform. Let _*g*_ denote the set of isoforms associated with gene *Ig*. The model contains a gene-level prediction head and an isoform-level prediction head:

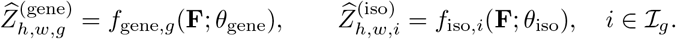

Non-negative output activations are used so that both predictions can be interpreted as pixel-level molecular abundance. Their corresponding spot-level predictions are

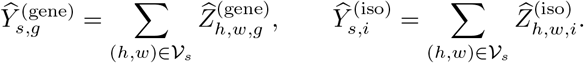

The supervised reconstruction objective is defined as

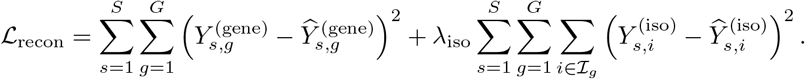

#### Gene–isoform consistency regularization

The total expression of a gene should be consistent with the combined abundance of its constituent isoforms. We therefore introduce a pixel-level consistency loss:

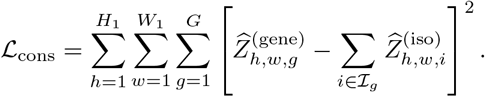

Unlike a consistency constraint imposed only after spot aggregation, this objective directly regularizes the latent allocation of gene expression among isoforms at every pixel. The complete training objective is

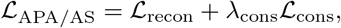

where *λ*_iso_ balances gene- and isoform-level reconstruction and *λ*_cons_ determines the strength of gene–isoform consistency. All model components are optimized jointly.

#### Pixel-level differential usage reconstruction

After training, the usage of isoform *i ∈ I*_*g*_ at pixel (*h, w*) is calculated as

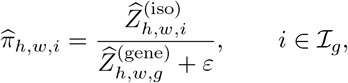

where *ε >* 0 is a small constant introduced for numerical stability. Equivalently, the denominator may be evaluated using the sum of the predicted isoform abundances:

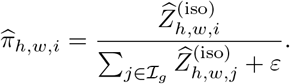

The gene–isoform consistency loss makes these two definitions approximately equivalent while separating changes in isoform usage from changes in total gene abundance. For two biological conditions or spatial regions *a* and *b*, pixel-or region-level differential usage is quantified by

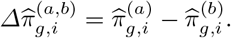

Consequently, the enhanced maps preserve the compositional relationship between gene-and isoform-level expression and support reconstruction of spatially localized differential APA or AS usage at pixel resolution.

### 4.7 Benchmark dataset generation

To quantitatively evaluate the accuracy of super-resolved expression prediction, we generated pseudo-spot datasets from high resolution spatial data with single- or subcellular resolution.

For spatial transcriptomics data, we generated benchmark datasets from Xenium, Xenium Prime, Stereo-seq and Visium HD data. Xenium is a targeted spatial transcriptomics platform that profiles predefined gene panels at subcellular resolution (approximately 0.2 *×* 0.2 *µ*m^2^) using in situ RNA detection. Xenium Prime provides spatial gene expression measurement at the same resolution as Xenium but with a larger panel of genes. Stereo-seq is a high-resolution spatial transcriptomics platform that captures whole-transcriptome RNA on DNA nanoball arrays at subcellular-scale resolution (0.5 *×*0.5 *µ*m^2^). Visium HD is a whole-transcriptome spatial profiling platform that captures RNA on a high-density array of 2 *×* 2 *µ*m^2^barcoded squares. Spot-level pseudo-Visium inputs were constructed by aggregating high-resolution pixel-level counts into spots that replicate the Visium array geometry. In particular, transcript counts were summed within a hexagonal grid of circular spots, each with a diameter of 55 *µ*m and spaced 100 *µ*m center-to-center, mirroring the layout of the Visium capture array.

For spatial proteomics data, we adopted subcellular resolution platforms to obtain ground truth measurement and generate pseudo-spot level data. Singular Genomics G4X is a subcellular-resolution spatial multiomics platform that simultaneously profiles targeted RNA and proteins using in situ sequencing and oligo-conjugated antibodies. Current G4X configurations support roughly 500-plex RNA and 18-plex protein profiling. Xenium also enables simultaneous spatial profiling of targeted RNA and protein markers at subcellular resolution using in situ RNA detection and oligonucleotide-conjugated antibodies, enabling up to 500 RNA targets and 27 proteins. As mentioned before, high-resolution data were aggregated to generate pseudo-spot level data for enhancement and analysis.

For spatial ATAC-seq data, we found limited datasets paired with high-resolution H&E image, which is required for our algorithm. Thus we adopted a recently published method called ISON to generate ATAC data from RNA measurements. We applied ISON on a Human Tonsil Visium HD data. For robustness, we aggregated the original 8 *×*8*µm*^2^measurements to 32*×* 32*µm*^2^, and used ISON to predict ATAC data at peak level, using a human tonsil single cell multiome data as reference. We then used the 32 *×* 32*µm*^2^level RNA and ATAC data as ground truth and aggregated them to a 64*×*64*µm*^2^grid for downstream enhancement and analysis.

### 4.8 Tissue segmentation with BANKSY

Tissue segmentation was performed on both reconstructed and ground-truth expression profiles using BANKSY. The original data were represented on a regular spatial grid at 8 *µ*m *×*8 *µ*m resolution. Prior to segmentation, each 2 *×*2 block of adjacent pixels was aggregated to generate expression profiles at 16 *µ*m*×* 16 *µ*m resolution. This aggregation reduced expression sparsity and improved the robustness of spatial domain identification. The same procedure was applied to reconstructed and ground-truth data to ensure directly comparable inputs.

The aggregated expression matrices and corresponding spatial coordinates were then analyzed with BANKSY using the default settings recommended for Visium HD-like data in the BANKSY tutorial. BANKSY-derived cluster assignments were used as tissue segmentation labels for downstream comparison of reconstructed and ground-truth spatial expression patterns.

### 4.9 Cell type deconvolution and annotation

Cell-type annotation of reconstructed and ground-truth spatial expression profiles was performed using RCTD. The original expression data were represented on a regular spatial grid at 8 *µ*m *×*8 *µ*m resolution. Based on evaluation across different spatial resolutions, aggregation to 16*×* 16 *µ*m^2^yielded more robust cell-type assignments. We therefore aggregated each 2*×* 2 block of adjacent pixels prior to RCTD analysis. The same aggregation procedure was applied to reconstructed and ground-truth expression profiles to ensure directly comparable inputs.

RCTD was subsequently applied to the 16*×* 16 *µ*m^2^-resolution expression profiles using the default settings recommended for Visium HD format data and cell-type annotation in the RCTD documentation. Cell-type identities inferred by RCTD were used for downstream comparison of cellular organization between reconstructed and ground-truth spatial expression maps. RCTD-derived cell-type proportions used as input to the reconstruction model were generated separately as described in the model workflow.

### 4.10 Evaluation metrics for expression prediction

Reconstruction accuracy was evaluated using root mean squared error (RMSE) and the structural similarity index measure (SSIM), which provide complementary assessments of quantitative reconstruction accuracy and spatial pattern preservation. RMSE measures point-wise differences between reconstructed and ground-truth expression values, with lower values indicating greater quantitative agreement. SSIM evaluates the similarity of spatial structures by considering local intensity, contrast, and spatial organization, with higher values indicating better preservation of spatial expression patterns.

To reduce the influence of extreme expression values, values above the 99.99th percentile were clipped to the corresponding percentile threshold before evaluation. Following clipping, expression values were rescaled using min-max normalization. For each comparison, the same normalization range was applied to all samples being compared to ensure that reconstructed and ground-truth expression maps were evaluated on a consistent scale. RMSE and SSIM were then calculated between each normalized reconstructed expression map and its corresponding ground-truth map.

## 5 Data availability

We analyzed the following publicly available datasets: (1) 10x Xenium human breast cancer data (https://www.10xgenomics.com/products/xenium-in-situ/preview-dataset-human-breast);(2) 10x Xenium Prime 5K mouse brain hemisphere data (https://www.10xgenomics.com/datasets/xenium-prime-fresh-frozen-mouse-brain); (3) 10x Xenium Prime 5K human breast cancer data (https://www.10xgenomics.com/datasets/xenium-prime-ffpe-human-breast-cancer); (4) 10x Xenium Prime 5K human lymph node data (https://www.10xgenomics.com/datasets/preview-data-xenium-prime-gene-expression); (5) 10x Xenium Prime 5K human ovarian cancer data (https://www.10xgenomics.com/datasets/xenium-prime-ffpe-human-ovarian-cancer); (6) 10x Xenium human ovarian cancer data (https://www.10xgenomics.com/datasets/ffpe-human-ovarian-cancer-data-with-human-immuno-oncology-profiling-panel-and-custom-add-on-1-standard); (7) 10x Xenium human glioblastoma data (https://www.10xgenomics.com/datasets/ffpe-human-brain-cancer-data-with-human-immuno-oncology-profiling-panel-and-custom-add-on-1-standard); (8) 10x Xenium human lung cancer data (https://www.10xgenomics.com/datasets/ffpe-human-lung-cancer-data-with-human-immuno-oncology-profiling-panel-and-custom-add-on-1-standard); (9) 10x Xenium human pancreas data (https://www.10xgenomics.com/datasets/ffpe-human-pancreas-with-xenium-multimodal-cell-segmentation-1-standard); (10) 10x Xenium human gastric cancer data reported in Schroeder et al. (https://www.nature.com/articles/s41592-025-02770-8); (11) Stereo-seq mouse brain data (https://en.stomics.tech/col1241/index.html); (12) 10x Xenium human colorectal cancer data (https://www.10xgenomics.com/platforms/visium/product-family/dataset-human-crc); (13) 10x Visium human tonsil data (https://www.10xgenomics.com/datasets/visium-cytassist-gene-and-protein-expression-library-of-human-tonsil-with-add-on-antibodies-h-e-6-5-mm-ffpe-2-standard); (14) 10x Xenium human renal cancer data (gene and protein) (https://www.10xgenomics.com/datasets/xenium-protein-ffpe-human-renal-ccrcc); (15) Mouse brain transcriptomics and metabolomics co-profiling data (https://www.nature.com/articles/s41587-023-01937-y); (16) 10x Visium and SiT mouse brain spatial long read sequencing data (https://doi.org/10.1093/nar/gkad169); (17) 10x Visium HD human tonsil data (https://www.10xgenomics.com/datasets/visium-hd-cytassist-gene-expression-human-tonsil-fresh-frozen); (18) epi-Patho-DBiT spatial-FFPE-ATAC data of MALT (https://www.nature.com/articles/s41467-026-71576-9).

## 6 Code availability

The spEnhance Python package is available at https://github.com/dsong-lab/spEnhance.

## Acknowledgements

The authors appreciate the comments and feedback from members of the DS Lab at UConn Health (https://dsong-lab.github.io/).

