## Supplementary Material for "Trustworthy super-resolution reconstruction across spatial omics modalities"

### S1 Supplementary Figures

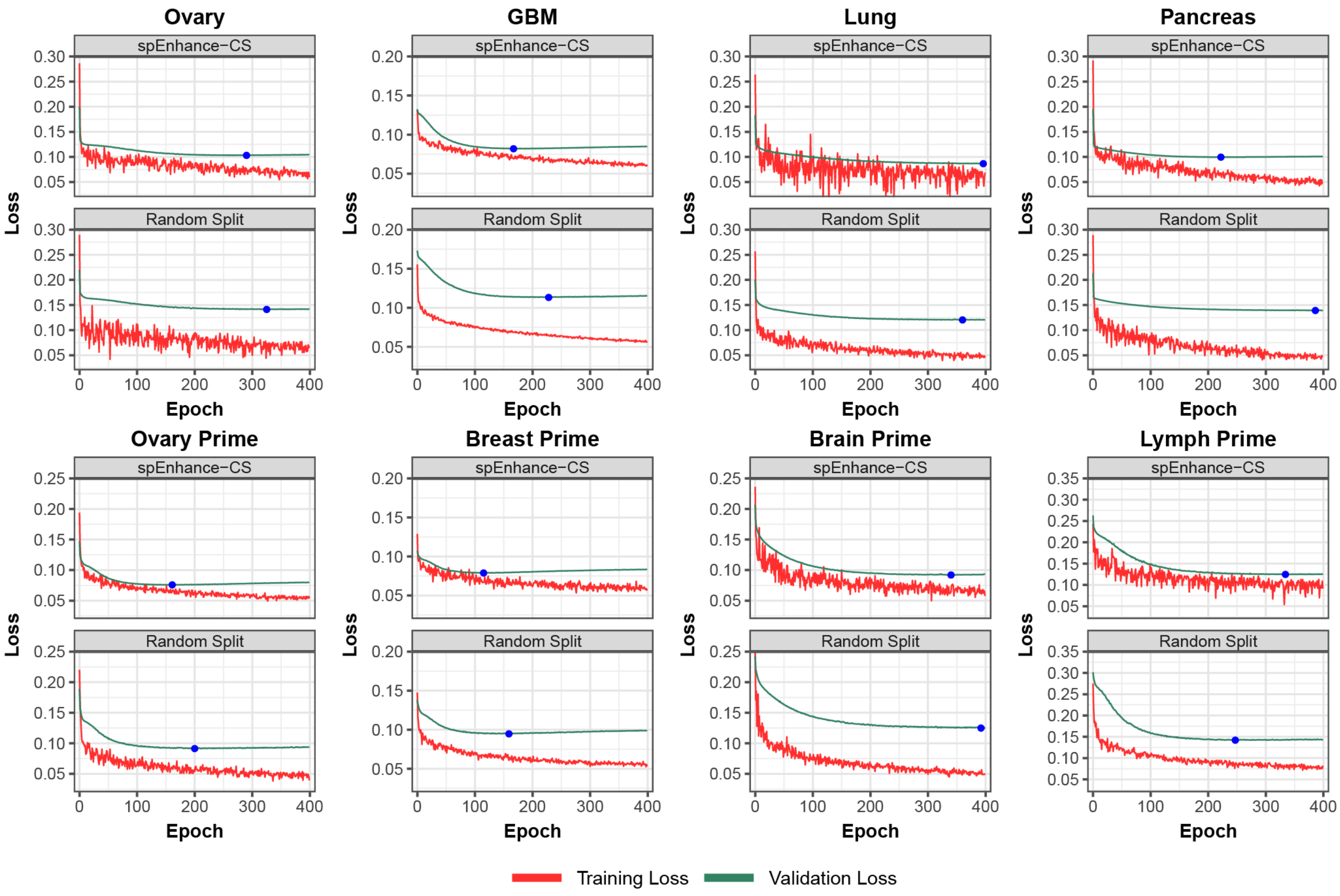


Fig. S1: Training and validation loss curves across epochs for spEnhance-CS and a random-split validation strategy in eight datasets. In each panel, the red curve shows training loss and the green curve shows validation loss; the blue point marks the selected epoch. Compared with random splitting, spEnhance-CS yields validation loss curves that more closely track the training loss trajectory while remaining based on statistically independent count partitions. This pattern suggests that spEnhance-CS provides a more reliable validation framework for model selection and reduces information leakage during training.

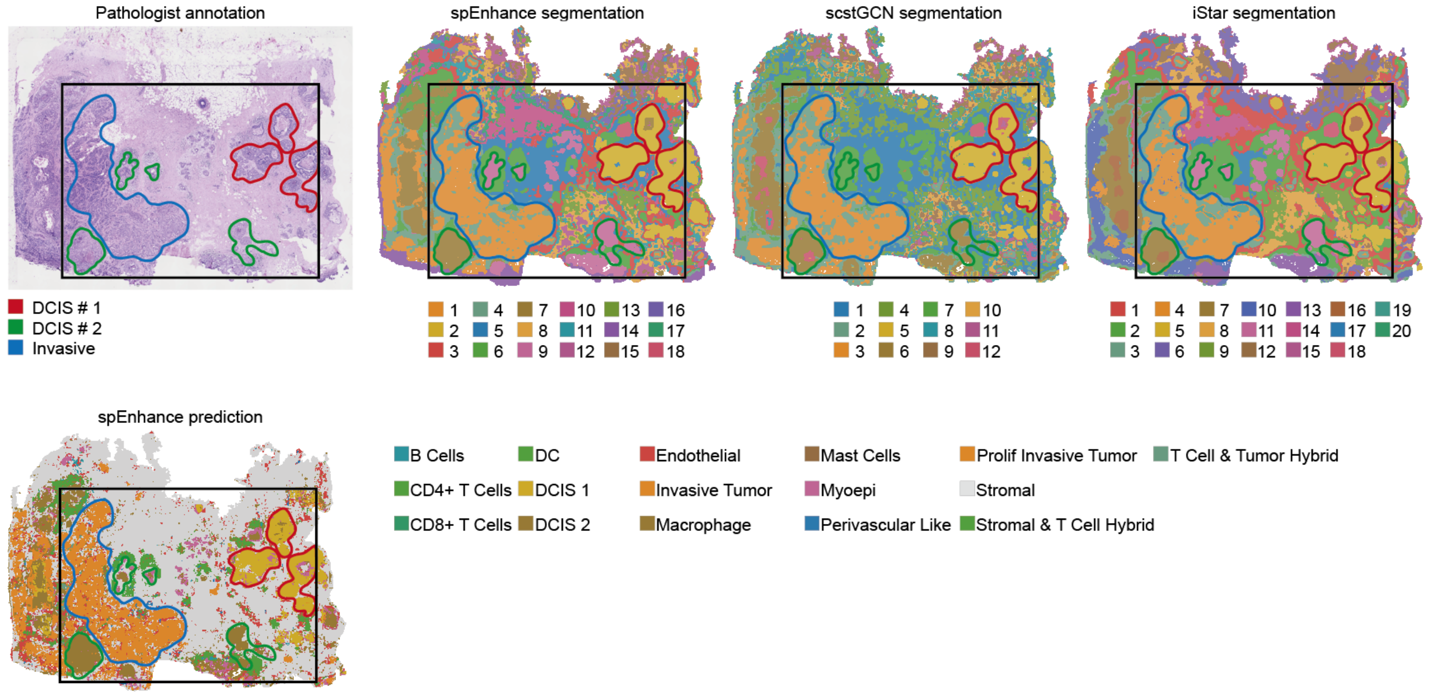


Fig. S2: Tissue segmentation of human breast cancer dataset. Visium measurement of the section was aligned to H&E image paired with Xenium data. Then high resolution spatial gene expression was reconstructed using spEnhance, scstGCN and iStar. BANKSY was used for tissue clustering and segmentation based on previous enhancement result. Pathologist annotation was adapted from [1]. spEnhance initial cell type prediction result was also shown for comparison.

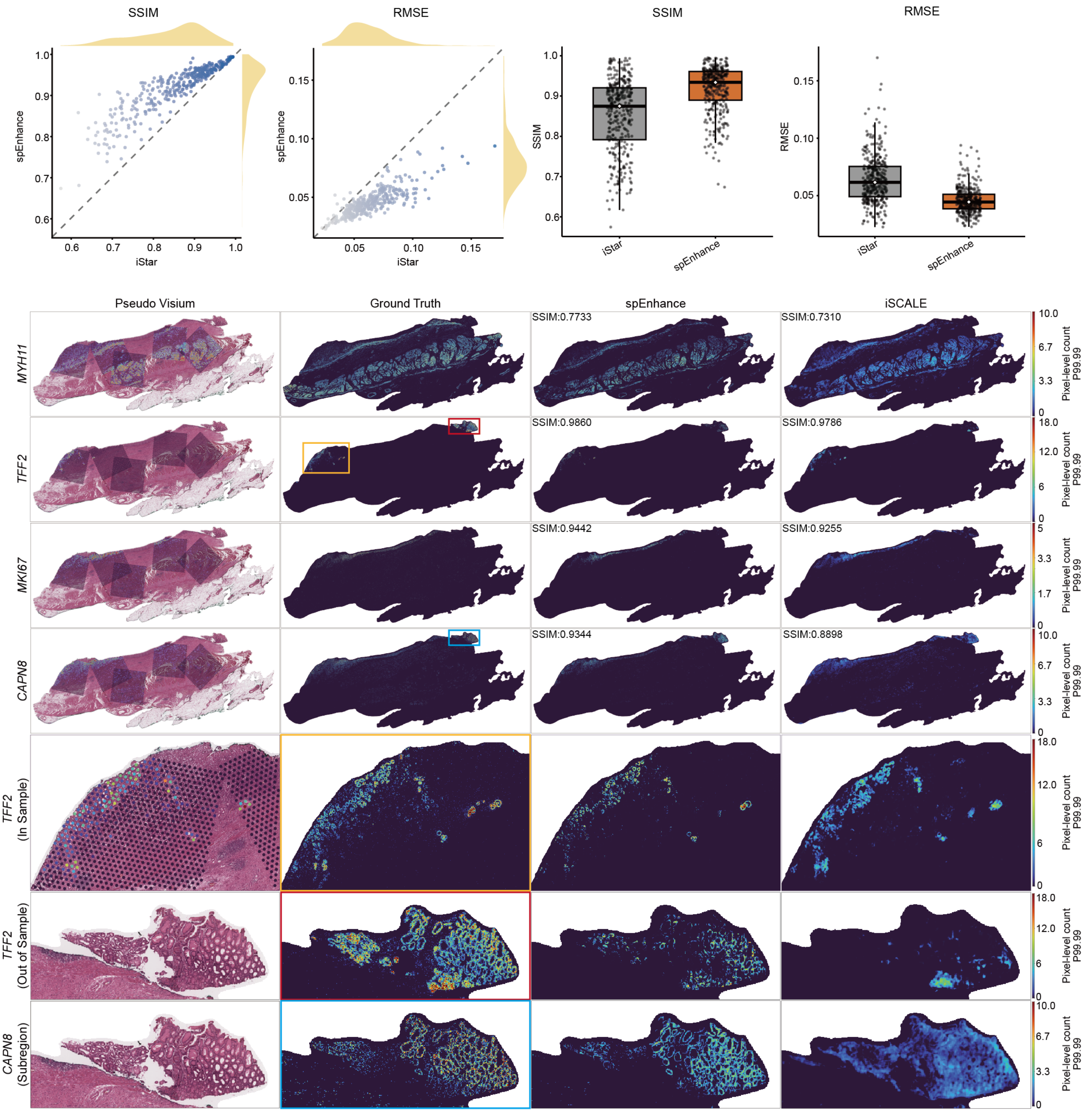


Fig. S3: Benchmark of spEnhance performance on multi-capture experiment design against iSCALE. Pseudo-Visium measurement of a human gastric cancer data from [2] was adopted. We used spEnhance and iSCALE for reconstructing high resolution spatial expression respectively. Scatter plots and boxplots comparing SSIM and RMSE of spEnhance and iSCALE compared with Xenium ground truth are shown. Zoom-in views of representative subregions are shown for visual comparison.

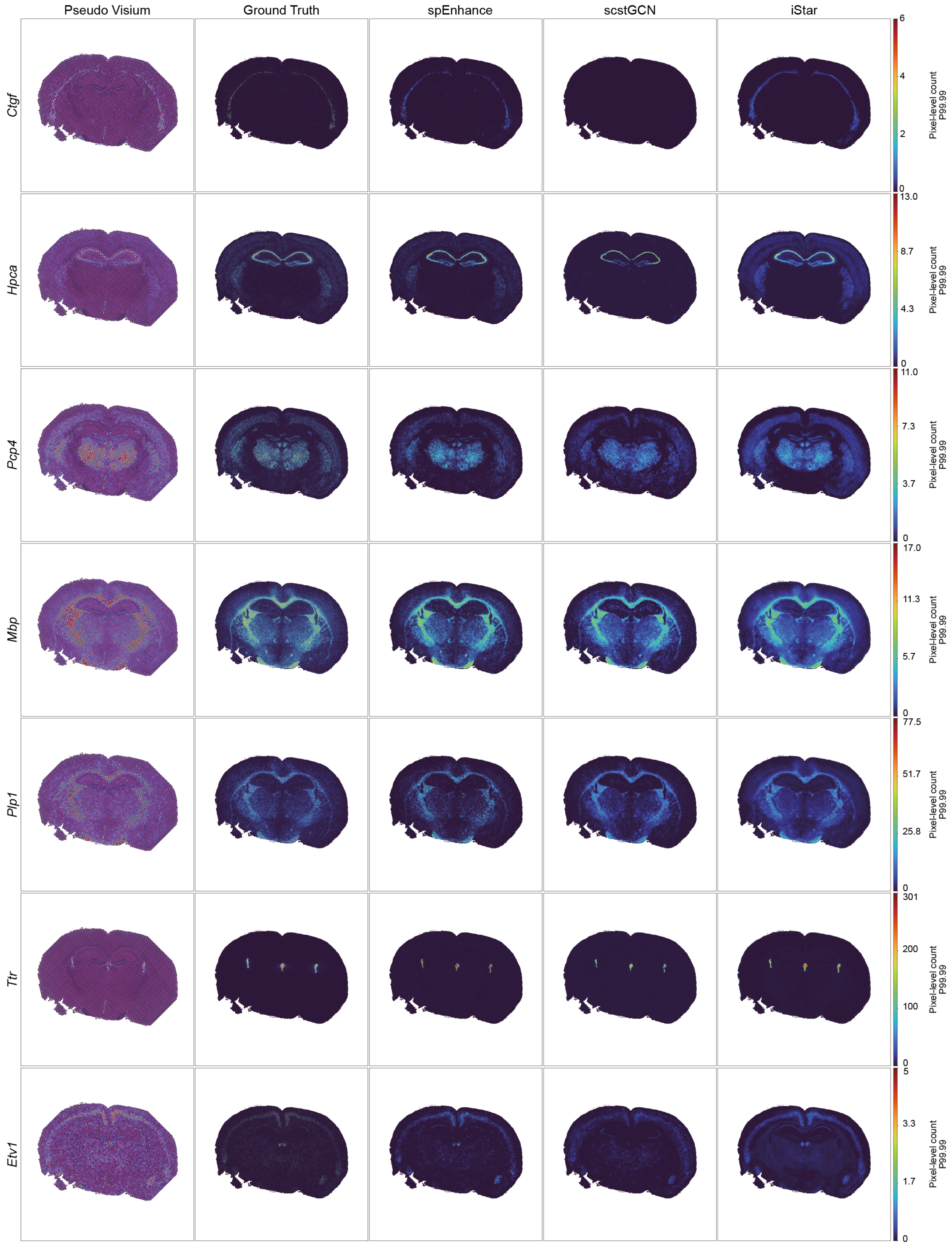


Fig. S4: Enhancement results of a mouse brain data. Pseudo-Visium data was generated from Stereo-seq measurements. spEnhance, scstGCN, iStar were used for high-resolution expression enhancement. Pseudo Visium data, Stereo-seq ground truth, and representative images of enhancement results from spEnhance, scstGCN and iStar were shown for visual comparison.
